# Towards whole-genome inference of polygenic scores with fast and memory-efficient algorithms

**DOI:** 10.1101/2025.01.19.633789

**Authors:** Shadi Zabad, Chirayu Anant Haryan, Simon Gravel, Sanchit Misra, Yue Li

## Abstract

With improved Whole Genome Sequencing (WGS) and variant imputation techniques, modern Genome-wide Association Studies (GWASs) have enriched our understanding of the landscape of genetic associations for thousands of disease phenotypes. However, translating the marginal associations for millions of genetic variants to integrated polygenic risk scores (PRS) that capture their joint effects on the phenotype remains a major challenge. Due to technical and statistical constraints, commonly-used PRS methods in this setting either perform heuristic Pruning-and-Thresholding or overlook most genetic association signals by restricting inference to small variant sets, such as HapMap3. Here, we present a set of algorithmic improvements and compact data structures that enable scaling summary statistics-based PRS inference to tens of millions of variants while avoiding numerical instabilities common in such high-dimensional settings. These enhancements consist of highly compressed Linkage-Disequilibrium (LD) matrix format, which integrates with streamlined and parallel coordinate ascent updating schemes. When incorporated into our existing PRS method (`VIPRS`), the new algorithms yield over 50 fold reductions in storage requirements and lead to orders of magnitude improvements in runtime and memory efficiency. The updated `VIPRS` software can now perform Variational Bayesian regression over 1.1 million HapMap3 variants in under a minute. Using this new scalable implementation, we applied `VIPRS` to 75 of the most heritable, continuous phenotypes in the UK Biobank, leveraging marginal associations for up to 18 million bi-allelic variants. Performing inference over this rich association data requires less than 20 minutes of wallclock time and 15GB of memory per phenotype. It also delivers consistent gains in cross-population transferability, with an average improvement of 10-15% in incremental R-squared.

## 1 Introduction

Polygenic risk scores (PRS) have recently garnered widespread interest in the clinical research community as a promising tool for patient stratification and personalized medicine [1–3]. Polygenic scores quantify the genetic component of complex traits and their inference can be framed as a high-dimensional multiple regression problem mapping from genotype to the phenotype of interest [4]. Many methods have been proposed to infer PRS from large-scale genome-wide association studies (GWASs), each of them tailored to deal with the unique challenges inherent in this form of data (see [5–7] and references therein). Because individual-level data is rarely publicly available due to privacy concerns, many PRS inference methods rely on GWAS summary statistics, matched with an appropriate Linkage-Disequilibrium (LD) reference panel [5, 6, 8–13].

While summary statistics-based PRS methods have been successfully deployed and tested in a variety of biobanks and research settings [14], they still face unique challenges of their own. First, these methods often require computing and publicly disseminating large-scale LD matrices from various reference panels to enable users to perform inference on their own GWAS datasets. LD matrices published by popular PRS methods, even when restricted to a relatively small subset of variants and represented in sparse and banded forms, require several Gigabytes of storage [10–12, 15]. Anything beyond this restricted variant set requires tens of Gigabytes to Terabytes of storage (cf. [15]), which makes inference and dissemination difficult. The second challenge concerns heterogeneities between the GWAS summary data and LD reference panels, including differences in allele frequency or LD patterns. Previous work has highlighted the many issues that can arise as a result of these differences, from numerical instabilities to poor prediction accuracy [13, 15, 16]. Finally, as better variant imputation tools and Whole Genome Sequencing (WGS) technologies become increasingly integrated into GWAS pipelines, performing PRS inference over millions of variants remains a technical challenge. Commonly-used PRS methods usually require hours of runtime and tens of Gigabytes of memory for a single phenotype, even when the regression is restricted to a small subset of variants [6, 15]. Scaling these methods to larger variant sets usually requires expensive specialized hardware.

Here, we present a major methodological update to our previously published work on Variational Inference of Polygenic Risk Scores (`VIPRS`) [13] and provide generally applicable solutions to these challenges. These solutions are summarized in the following four contributions. First, our new software tools provide utilities to compute large-scale LD matrices and compress them by over 50 fold. Using our new LD matrix storage format, we show that LD matrices for 1.4 million HapMap3+ variants can be shrunk to as little as 300MB, making them easily shareable via typical workplace communication channels. Second, our study provides concrete recommendations for estimating and constructing well-conditioned LD matrices that behave stably in ultra high dimensions. Third, we implemented and tested a new low-memory version of the coordinate ascent variational inference (CAVI) algorithm that operates directly on the compressed LD data, which reduces memory footprint by more than order of magnitude. Finally, we upgraded our software tools to use two layers of parallelism, within coordinate ascent and across independent chromosomes, which provides substantial speedups in inference time, up to two orders of magnitude faster than previously published versions of the software [13]. Putting all of this together makes `VIPRS` one of the most computationally efficient methods for model-based PRS inference, rivaling even highly optimized implementations of heuristic methods such as Clumping-and-Thresholding (C+T) [9, 13, 17].

To demonstrate the promise of these methodological improvements, we applied the latest version of the `VIPRS` software to 75 of the most heritable phenotypes in the Pan-UKB resource [18] using all well-imputed bi-allelic genetic variants, reaching up to 18 million variants in European samples. Our analyses demonstrate that the updated variational Bayesian algorithm is able to converge on data of this scale in less 20 minutes, using less than 15 Gigabytes of RAM. In cross-population validation analyses, we show that expanding the variant set used in the regression results in considerable improvements in transferability across most phenotype categories.

## 2 Material and Methods

### 2.1 Algorithms and data structures for scalable PRS inference

#### 2.1.1 Efficient Linkage-Disequilibrium (LD) matrix storage format and specification

One of the main computational bottlenecks for model-based PRS inference methods is the LD matrix and how it is represented, stored, processed, and ultimately deployed during inference. Recent work has highlighted the potential for orders-of-magnitude reductions in runtime and memory utilization for many routine computational tasks in statistical genetics when the LD martix (or its inverse) is highly sparse and compactly represented [19]. As a practical matter, this bottleneck has largely restricted many commonly-used PRS methods to a relatively small subset of roughly 1 million HapMap3 variants [10–12]. Recent work has explored various strategies and techniques to bypass the difficulty of working with large-scale LD matrices, either by transforming and sparsifying the LD matrix itself [15, 19] or developing highly scalable algorithms and heuristics that can still work with matrices at the scale of 7-10 million variants [13, 20]. While the latter approach is useful in principle, it is less scalable and moving or sharing data of this size can be cumbersome.

In this work, we sought to substantially reduce LD storage requirements and its associated costs in terms of memory usage and network traffic. To that end, we took inspiration from LD matrix storage solutions provided by `hail` [21] (`BlockMatrix`) and the `bcor` storage format implemented in `LDStore` [22]. As summarized in **Figure** 1, the update encompasses three steps that are simple to implement, fast to run, and generalizable. These updates apply to all LD matrix estimators outlined in our previous work [13], including banded (windowed), block-diagonal, and shrunk LD matrices.

**Figure 1:**
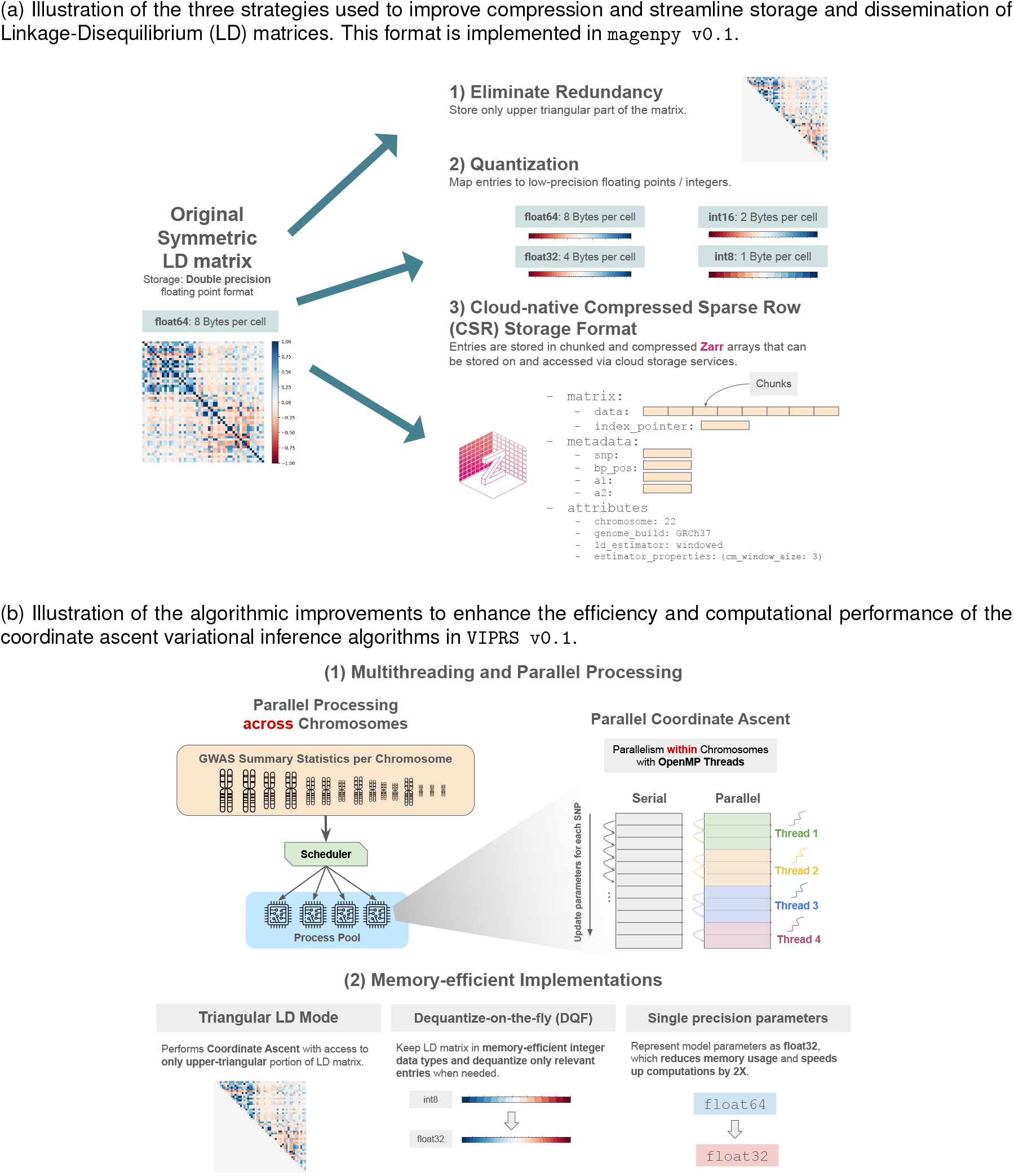
Diagrams illustrating algorithmic and data structure improvements in the latest version (`v0.1`) of the Variational Inference of Polygenic Risk Scores (`VIPRS`) software.

The first step eliminates redundancy and only stores the upper-triangular portion of the symmetric LD matrix (not including the diagonal itself). This step reduces storage requirements by more than a factor of two. The second step involves mapping the entries of the LD matrix to lower-precision floating points or quantizing to integer data types. Previous work has shown that rounding the entries of the LD matrix up to two decimal points has minimal impact on heritability estimation methods [23]. In the context of fine-mapping, the `LDStore` software [22] for computing and storing LD matrices (developed in conjunction with the `FINEMAP` method [24]) has explored similar compression mechanisms, though documentation of the precise techniques and their impact on inference quality is lacking. We are not aware of similar work in the context of polygenic risk score inference. Mapping to lower-precision floating points is as simple as casting the entries of the LD matrix from double precision floating points (`float64`) to single precision floating point format (`float32`). This alone can reduce the storage requirements by a factor of 2, going from 8 to 4 bytes per entry. In principle, we can shrink the storage requirements even further by going to half precision floats (`float16`). However, we didn’t explore the half precision encoding of the data because there is not a universal standard for encoding this data type across programming languages and platforms. Instead, we explored quantizing the data to low-precision integers as an alternative, which allowed us to encode the entries of the LD matrix using a single byte (`int8`) or two bytes (`int16`) per entry, representing a reduction of up to a factor of 8 in storage requirements. **Table** 1 summarizes the implemented data types, their storage requirements, and resolution. Quantization is done using the “scale quantization” technique [25] where the quantized entry 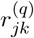 is computed as:

**Table 1:** Data types for representing the entries of the LD matrix and their properties.

| Data Type | Number of bytes | Resolution |
| --- | --- | --- |
| float64 | 8 | $10^{-15}$ |
| float32 | 4 | $10^{-6}$ |
| int16 | 2 | $2^{-15} \approx 0.00003$ |
| int8 | 1 | $2^{-7} \approx 0.008$ |

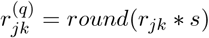

where *r*_*jk*_ is the Pearson correlation coefficient between variants *j* and *k*, the scaling factor *s* = 2^*b*−1^ is the maximum representable positive value for the integer data type, *b* is the number of bits used to encode the integer, and *round* maps the product to the nearest integer. Dequantization reverses the scaling operation: 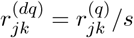.

Finally, to support the new optimized implementations of the coordinate ascent algorithm in `VIPRS`, we updated the storage format from Variable Length (`VarLen`) Arrays to Compressed Sparse Row (CSR) format. In this new format, the non-zero entries of the LD matrix are stored contiguously in a very long one-dimensional (1D) array. To delineate the boundaries of each row, we also store an “index-pointer” array that records the start index of each row (**Figure** 1). This data is stored in a hierarchical, compressed, chunked, and cloud-native storage format known as `Zarr` [26], which supports multi-threaded read and write access. In addition to the entries of the LD matrix itself, the hierarchical `Zarr` store includes important metadata about the variants represented in the matrix, such as variant rsID, position, reference and alternative allele, LD score, etc. To encourage reproducibility, various “attributes” about the LD matrix may be added to the `Zarr` hierarchy, including the Biobank from which the matrix was computed, the ancestry of the samples, the sample size, the LD estimator and its properties, etc. This structure is summarized in **Figure** 1.

Other features of the `Zarr` format are also important to highlight in this context. For instance, flexible and powerful compression API can help shrink the size of the stored data even further. In the latest release, we now use the `Zstandard` compression algorithm by default, instead of the `lz4` compressor used in the old format. For LD data, the `Zstandard` compressor can provide a compression ratio of up to 4 (data not shown). It is also worth emphasizing that `Zarr` is cloud-native storage format [26], and this opens the way to provide read access to relevant portions of large-scale LD matrices from cloud storage services directly, without the need to download them locally. Finally, the `Zarr` storage format is supported through specialized APIs in many popular programming languages, allowing for standardized and universal LD storage specification that can be used for downstream analyses across many statistical genetics toolkits.

#### 2.1.2 Optimized and memory-efficient coordinate ascent algorithms

Most model-based PRS inference methods follow iterative, coordinate-wise sampling or optimization techniques, where the computational cost is dominated by cycling through millions of variants and sampling/updating their parameters conditioned on the parameter values of all the other variants [8, 11–13, 27]. In the case of `VIPRS`, this is represented by the E-Step of the Variational EM scheme, where we run a coordinate ascent algorithm to update the variational parameters associated with each variant [13]. According to our benchmarks, this step can account for up to 80% of the total runtime of the program (data not shown). Thus, optimizing coordinate-wise updates is essential for scaling PRS inference to tens of millions of variants. In the case of `VIPRS`, these optimizations consisted of five steps.

First, in earlier versions of the `VIPRS` software (`v0.0.4`), we found that the coordinate ascent step was slowed down due to the `VarLen` array encoding from `Zarr` necessitating interaction with the `python` interface, where loops over millions of elements are relatively inefficient. In the new software implementation, we re-wrote the coordinate ascent step in pure `C/C++`, which necessitated using the Compressed Sparse Row (CSR) format for the LD matrix described previously. This update alone resulted in at least 10-fold improvements in speed over the older version of the software (**Supplementary Figure** S1).

Second, we optimized the linear algebra operations in the coordinate ascent updates. If a `BLAS` library [28] was detected on the user’s system during installation, we delegated the intensive linear algebra computations to these optimized subroutines for best performance. The manually-written linear algebra functions, to which the program reverts if BLAS is not available, were also annotated with compiler hints to use Single Instruction/Multiple Data (SIMD) intrinsics where possible, which should enhance their performance considerably.

Third, through profiling, we found that the coordinate ascent step had become memory bandwidth bound after the above improvements. Therefore, we explored casting all the parameter data types as well as the input data to single precision floating point (`float32`) format instead of double precision (`float64`), which reduced memory bandwidth pressure and total runtime by a factor of two (**Supplementary Figure** S1), without significantly affecting prediction accuracy (**Figure** 2). The latest version of `VIPRS` uses single precision floats by default, though the user has the option to request the inference be done in double precision.

**Figure 2:**
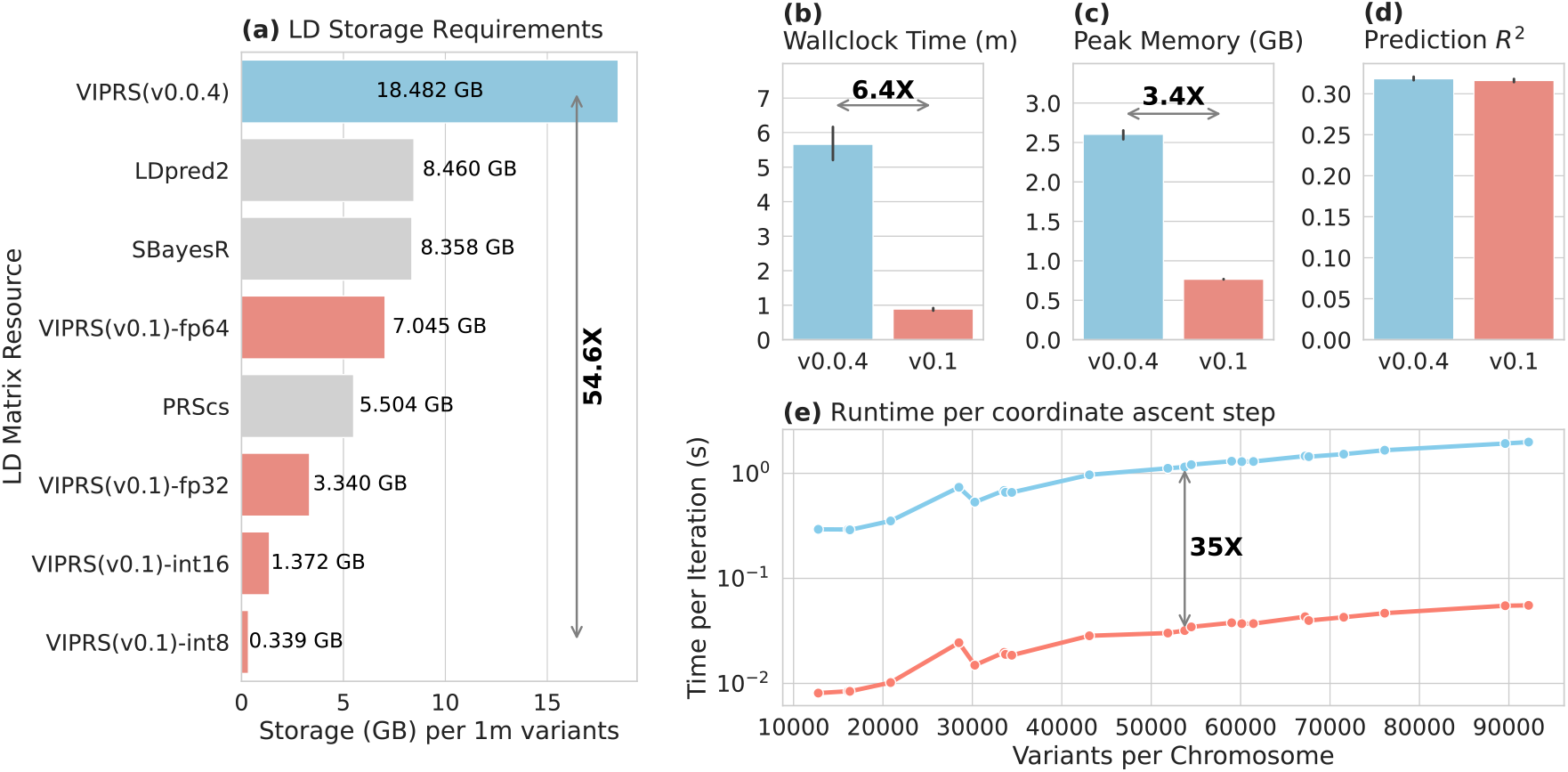
Comparing the computational performance and resource utilization of the old (`v0.0.4` ; skyblue color) versus newer (`v0.1` ; salmon color) versions of `VIPRS`, both with their default settings. The two versions are benchmarked on GWAS summary statistics for Standing Height from the UK Biobank, with ≈ 1.1 million HapMap3 variants included in the analysis. Panel **(a)** shows the storage requirements for Linkage-Disequilibrium (LD) matrices, with matrices from commonly-used PRS methods (grey bars) included for comparison. `VIPRS v0.1` supports various data types for storing the entries of the LD matrix, including two float precisions (`float64` and `float32`) and, with quantization, two integer data types (`int16` and `int8`). **(b)** and **(c)** show the total wallclock time (minutes) and peak memory usage (GB) of the two versions of `VIPRS`. Panel **(d)** shows the prediction accuracy (R-squared) on held-out test sets in the UK Biobank. Panel **(e)** shows the runtime per iteration (in seconds and on log-scale) of the two versions of the software as a function of the number of variants on each chromosome. Arrows and bold text in panels **(a-c)** and **(e)** indicate the mean fold improvement of the new software version over the old. Vertical black lines above the bars in panels **(b-d)** show standard errors of each metric across 5 folds of the data.

Fourth, in our previous work, we showed that the variational coordinate ascent updates can be written so that we access and multiply with the symmetric LD matrix only once per iteration (Supplementary Note of [13]; Algorithm 1). In our latest implementation, we provide the option to perform these updates with access to only the upper-triangular portion of the LD matrix (Algorithm 2), thus obviating the need to reconstruct the lower-triangular half and reducing memory usage significantly. To provide an intuition, it helps to think of the original “right-hand updating scheme” [11, 13] outlined in Algorithm 1 as a message-passing algorithm, where variant *j* updates its parameters and then broadcasts the information about the change to all its neighbors. In this paradigm, each variant stores information about their neighbors’ posterior mean (weighed by their LD) in the *q*-factor (*q*_*i*_:=Σ_*j*≠*i*_ *R*_*ij*_*η*_*j*_). Under the assumption that the variants are updated in order, it is easy to show that, at any given iteration, it is sufficient for the variant to broadcast the information to variants that come after it (line 7 of Algorithm 2). Then, after the coordinate ascent loop terminates, the information about the changes for variants that come after *j* can be propagated backwards with a simple sparse matrix-vector multiplication with the upper-triangular portion of the LD matrix *R*^(*UT*)^ (line 13 of Algorithm 2). This “low-memory” or “triangular LD” mode is now the default for the latest version of `VIPRS`.

##### Algorithm 1

Coordinate Ascent Variational Inference (**Symmetric LD**)

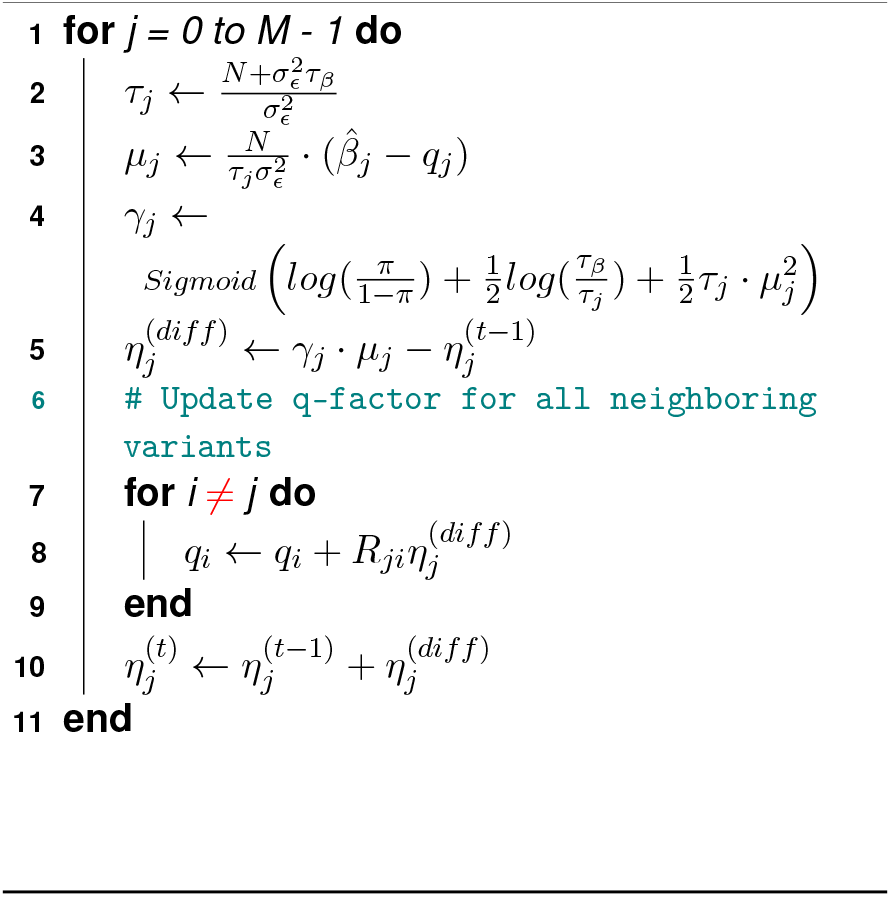

##### Algorithm 2

Coordinate Ascent Variational Inference (**Triangular LD**)

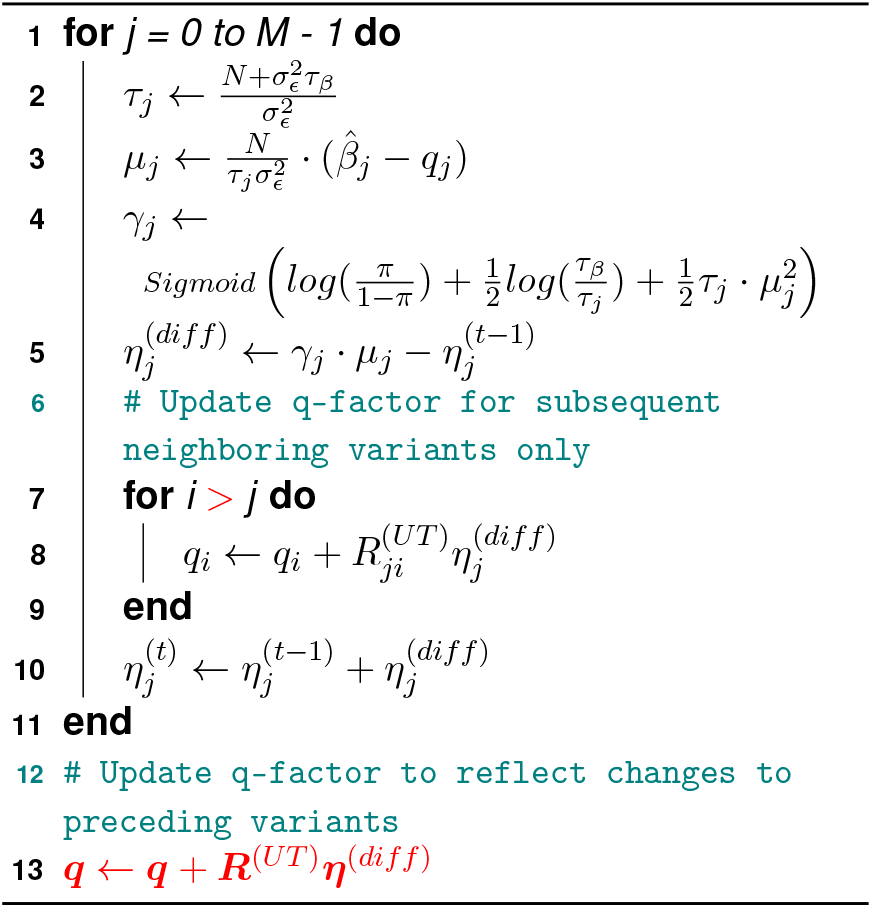

Finally, to reduce memory utilization even more, we provide the option for the user to run the software without dequantizing the entire LD matrix beforehand. In this framework, the relevant entries of the matrix are dequantized anew (hence `dequantize-on-the-fly` or DQF) whenever they are needed during the program’s runtime. This should reduce the memory usage from the LD matrix during inference by up to a factor of 8 (`int8` versus `float64` representation of LD data), though it comes at a minor cost of extra overhead and redundant computations.

#### 2.1.3 Taking advantage of multi-core systems: Parallel coordinate ascent and parallel processing across chromosomes

While the optimizations discussed in the previous section significantly boosted the performance of the software, the algorithm as outlined is still inherently serial and does not take full advantage of modern multi-core computing environments. Other PRS methods already take advantage of parallel processing capabilities in various ways, which enhanced their efficiency considerably [8, 11, 12, 20, 29]. To make strides in this direction, we identified two areas where parallelism across roughly independent tasks might prove useful for `VIPRS`.

In the first case, we explored performing inference in parallel across chromosomes, where the GWAS summary statistics and LD reference panel for each chromosome are read and analyzed independently by different system-level processes (**Figure** 1). This task is done with the help of a process pool interface provided by the `joblib` python library [30], where the user can specify the total number of processes that the system should run in parallel. One potential drawback of this approach is that it may result in extra overhead and increase memory usage, depending on the number of processes running concurrently. Nevertheless, it can speed up runtime significantly and it can be accessed by simply setting the flag `–-n-jobs` when running our updated command-line interface (CLI) script `viprs_fit` (see **Code Availability**).

In the second case, we explored performing parallel updates within the coordinate-ascent algorithm itself by leveraging the `OpenMP` multi-threading Application Programming Interface (API) [31]. Parallel coordinate ascent algorithms have been studied extensively in the literature, especially with regards to their convergence properties [32]. In `VIPRS`, the implementation of this scheme is as follows: Instead of updating the parameters of the variants serially as in Algorithm 1, we divide the variants into chunks and update these chunks simultaneously and in parallel with `OpenMP` threads (**Figure** 1). In idealized settings, this should speed up both Algorithms 1 and 2 by a factor proportional to the number of threads. However, in practice, there may be some overhead, especially for smaller chromosomes where the runtime-per-iteration is already negligible. An important detail to highlight here is that, unlike some previous implementations (e.g. `Lassosum` [8]), our parallel coordinate ascent algorithm is not restricted to LD blocks and works seamlessly with any LD matrix structure (windowed, shrunk, or block).

We caution users regarding multi-threading in the coordinate ascent step paired with Triangular LD as in Algorithm 2. In our experiments, we noticed that this setup, when coupled with a small variants/threads ratio, can result in some instabilities and oscillations in the parameter values. This is due to “staleness” of the parameters (e.g. *q*-factor is one iteration behind) and such effects have been documented in the literature on parallel and asynchronous coordinate ascent [33, 34]. To resolve this issue, we implemented a heuristic that decreases the number of threads by one when instabilities are detected. In future work, introducing a step size (or learning rate) as suggested by Liu et al. [34] may be a more appropriate and general solution. In addition, staleness and oscillations can make the convergence slower, which necessitates running the EM algorithm for more iterations, hence counteracting the benefits of multi-threading in these cases.

#### 2.1.4 General software usability improvements

The algorithmic improvements described earlier would be inaccessible without software design and tools that are easy to distribute, in addition to being well-tested and documented. Since their last release, our python packages `magenpy` (our in-house toolkit for GWAS and LD data harmonization and simulation) and `viprs` (see **Code Availability**) have undergone extensive refactoring to simplify the code, add basic unit and integration tests, make it more robust, and document most of the modules, functionalities, and interfaces. Specifically, we now follow a Continuous Integration/Continuous Development (CI/CD) testing regime where any updates to the software have to pass basic functionality testing across different platforms, including Windows, MacOS, and Linux. Furthermore, both packages now have dedicated web pages with documentation that include installation guidelines, walk-throughs of the basic features, basic tutorials, and API references. These resources are not yet complete and we anticipate that they will grow further in the future, especially as they gain interest from the research community. Additionally, there remains some challenges with distributing the software across platforms, given the wide-range of dependencies that the packages require. To facilitate distribution and widespread adoption, we also now provide `Docker` scripts [35] that containerize both `magenpy` and `viprs` in addition to all their essential dependencies. Both packages are open source and distributed under the MIT license.

### 2.2 Improving numerical stability of sumstats-based PRS methods

#### 2.2.1 Causes of numerical instabilities in PRS inference

In previous work, we noted that `VIPRS` and other summary statistics-based PRS methods are prone to numerical instabilities in some settings [13]. These numerical issues are characterized by the parameter estimates β exploding in magnitude and the Mean Squared Error (MSE) component of the objective function becoming negative. We initially hypothesized that these issues were partly caused by heterogeneities between the GWAS summary statistics and the LD reference panel and recommended performing Quality Control (QC) filtering using tools like DENTIST [36] before fitting the model. This explanation, however, was incomplete, as these numerical issues have been subsequently detected even in cases where the GWAS summary statistics and the LD panel originate from the same cohort.

Here, we propose that another main cause of these numerical pathologies can be traced back to the spectral properties of the LD matrix itself. Model-based PRS methods typically aim to optimize or approximate an objective that includes the MSE, a quantity that can be expressed in terms of the marginal effect sizes 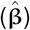 and the in-sample LD matrix (**R**):

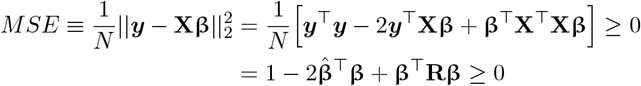

The equality holds under the assumptions that both the genotype matrix **X** and the phenotype vector ***y*** are fully observed, noise-free, and standardized column-wise. In practice, **X** can contain substantial amounts of missing values and noisy entries due to variant calling or imputation errors. Furthermore, even if **X** is perfectly observed, the entries of **R** are estimated in a pairwise fashion and then further shrunk, thresholded, and compressed. All of these sources of noise to can make our approximation of the in-sample LD, denoted as 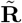, not positive semi-definite (PSD), which can lead the quadratic form 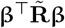 to become negative. To be precise, if the minimum eigenvalue of the approximate LD matrix 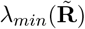 is strongly negative, the optimization for β can run away towards the corresponding eigenvector and create negative errors, leading directly to the numerical explosions reported in the PRS literature [13, 15].

Aside from the spectral properties of the approximate LD matrix 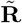, another problem that can lead to numerical instabilities is over-sparsifying or shrinking the LD matrix or substantial heterogeneities between the summary statistics and the LD reference panel [13, 15, 16]. This issue becomes especially prominent when the model requires estimating the residual variance 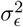. To take an extreme example, assume that we approximate **R** with the identity matrix **I**. While this choice is guaranteed to be PSD, it could nonetheless result in numerical instabilities because it will largely underestimate the quadratic form **β**^⊤^**Rβ**, which again can result in a negative MSE in some circumstances. This issue is harder to diagnose and satisfactorily address in practice, except via discarding heterogeneous summary data [16, 36].

#### 2.2.2 Analyzing the spectral properties of LD matrices

To facilitate analyzing the spectral properties of LD matrices, we developed data structures and dedicated software tools for computing extremal eigenvalues of LD matrices efficiently. Our software package `magenpy` now provides a linear operator class [37], called `LDLinearOperator`, that enables users to perform a wide range of linear algebra operations on LD matrices in their compressed form (i.e. while they remain triangular and quantized). To rapidly estimate the extremal eigenvalues of an LD matrix, we use iterative methods provided by `scipy` sparse matrix interfaces for the `ARPACK` library [37, 38]. These routines can estimate the minimum eigenvalue of chromosome-wide LD matrices with hundreds of thousands of variants in a matter of minutes, using limited memory footprint.

Our analyses indicate that many common approaches for estimating LD matrices in the context of PRS inference result in non-PSD matrices with large negative eigenvalues. Specifically, there are three main contributing factors that can impact the spectrum of estimated LD matrices: Sparsification pattern, pairwise correlation estimator in the presence of missing data, and approximation error. Sparsification pattern is the manner by which the sample LD matrix is transformed into a sparse one, using e.g. banded or block-diagonal masks. Our analyses indicate that banded LD matrices with fixed window size, e.g. 3 centiMorgan (cM), tend to have relatively large negative eigenvalues, especially when the sample size of the reference panel is small and/or in the presence of variants in long-range LD (LRLD) regions (**Supplementary Figure** S7). Block diagonal matrices are supposed to be more well-conditioned in principle, since each block is a Gram matrix by construction, and the eigenvalues of the entire LD matrix is simply the concatenation of the eigenvalues of individual blocks [39]. This observation explains why block-diagonal matrices are popular in PRS inference methods [8, 10], especially methods that are applied to larger variant sets [15].

However, even if block-diagonal LD matrices are used in practice, the resulting matrices may still have large negative eigenvalues because of the other two contributing factors. First, many pairwise correlation estimators from genotype data, such as `plink1.9` [17], use an unbiased estimator of Pearson correlation that discards missing observations. As a result, the entries of the LD matrix are estimated independently using disjoint subsets of the data. Thus, we expect that in the presence of missing genotype calls due to hard-thresholding, this will affect the spectra of estimated LD matrices [40]. Our analyses indicate that this indeed can result in matrices with large negative eigenvalues, even when using block-diagonal masks (**Supplementary Figure** S8). Instead of discarding missing genotypes, an alternative is to impute them using, e.g. Mean Imputation (MI) [40]. This approach produces LD matrices that are nearly positive semi-definite and have more stable spectrum (**Supplementary Figure** S8). For a detailed discussion of the benefits and drawbacks of both approaches, we refer the reader to the work of Choi and Tibshirani (2013) [40].

Finally, other sources of approximation error can also impact the spectra of estimated LD matrices. For example, thresholding individual entries or using ultra low-precision storage formats (e.g. quantization to `int8`) will negatively affect the spectrum, especially for large-scale LD matrices (**Supplementary Figure** S8). Therefore, there is a trade-off to be struck between compressibility and producing well-conditioned matrices.

#### 2.2.3 Proposed solutions and recommendations to achieve numerical stability

Although objective functions of the form discussed above have been carefully explored in the statistics literature for the past decade [40–42], it remains scarcely cited in statistical genetics, with some exceptions (e.g. [43]). In particular, the pioneering work of Loh and Wainwright (2012) [41] introduced losses of this form to allow for fitting penalized regression models when the design matrix **X** contains missing or noisy features. The authors noted the potential pitfalls with non-PSD covariance matrices and recommended constraining the **β** parameters during optimization to restore the convexity of the LASSO objective and to prevent numerical instabilities. In that original work, the authors proposed a projected gradient descent algorithm to perform this constrained minimization, which can be impractical in high dimensional settings. An alternative approach was proposed by Datta and Zou (2017) [42], where the authors advocated projecting 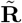 to the nearest PSD matrix and then using efficient coordinate ascent algorithms to optimize the modified objective. This approach is promising in principle, though projecting large-scale LD matrices onto the PSD cone can be prohibitively expensive and it may destroy the sparsity structure of 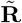. One way to bypass this difficulty is to work with block-diagonal LD matrices, where each block can be projected, denoised, and post-processed separately, as done in two recent PRS methods [15, 44] as well as in the software implementation of `PRScs` [10]. However, previous work indicated that banded LD matrices may perform slightly better than block-diagonal ones in some settings [13, 16], which motivated us to explore more general solutions.

Here, we followed a third approach outlined by Choi and Tibshirani (2013) [40], where instead of using the approximate LD matrix 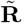 in the objective, we use 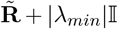, where |*λ*_*min*_| is the absolute value of the smallest (negative) eigenvalue of 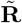 and 𝕀 is the identity matrix. This is similar in principle to the approach outlined by the developers of `Lassosum2` [16] method, though in that work the authors perform uninformed grid search over the hyperparameter instead of deriving it from the spectral properties of the LD matrix itself. In the case of `VIPRS`, this simple modification affects the Evidence Lower BOund (ELBO) [13], which includes the expectation of the MSE with respect to the variational density 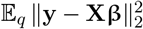. Building on our previous work [13] and replacing 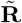 with 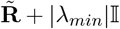, this expectation evaluates to:

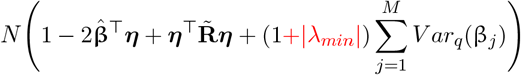

The change from the original expression is highlighted in red. This, in turn, affects the updates for the variational parameters, mainly through the posterior variance estimate 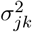:

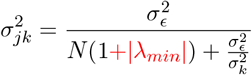

Thus, the main effect of adding the |*λ*_*min*_| to the objective is that it constrains the posterior for the effect size and it acts as an additional shrinkage penalty [16, 40]. The strength of this penalty is proportional to the magnitude of the smallest negative eigenvalue: The more ill-conditioned the LD matrix, the stronger the penalty. In practice, when working with large-scale banded LD matrices, we found that this can result in over-shrunk effect size estimates, which may lower prediction accuracy.

To deal with the potential for over-shrinkage and under-fitting, we provide two alternative solutions. In the first case, if independent validation dataset is available, we allow the user to explore an informed grid over multiples of the parameter |*λ*_*min*_|, e.g. the grid can be constructed as: {0, 0.01, 0.1, 1, 2} ∗ |*λ*_*min*_| [16, 40]. *Then we choose the penalty level that maximizes prediction on the held-out validation set*. In the second case, instead of using a single |*λ*_*min*_| penalty chromosome-wide, we allow the penalty to vary over genomic blocks. This is inspired by our observation that some genomic regions (e.g. long-range LD) contribute differentially to the negative eigenvalues of the LD matrix (**Supplementary Figure** S7), and penalizing variants in those problematic regions more strongly will stabilize the optimization algorithm without over-shrinking variants in more well-behaved regions.

In summary, our recommendations for constructing large-scale and well-conditioned LD matrices are as follows. In general, we recommend using block-diagonal masks for sparsifying the matrix. If the genotype matrix contains missing genotype calls, it is advisable to perform either Mean Imputation (MI) or estimate pairwise correlation based on raw dosages. To reduce approximation errors, it may be beneficial to use `int16` data type for storage, though in our experiments we found that `int8` works fine in most cases. If any of these conditions are unattainable for technical or other reasons, we recommend using the |λ_*min*_| penalty approach to constrain the parameter estimates and stabilize the inference process.

### 2.3 Benchmarking experiments

#### 2.3.1 Data used for benchmarking

To assess the impact of the algorithmic modifications introduced in the latest version of `VIPRS`, we used a subset of the summary statistics and LD reference data that was generated for the original `VIPRS` manuscript [13]. For the LD reference panel, we used LD matrices for approximately 1.1 million HapMap3 variants, computed using the windowed estimator with a window size of 3 centiMorgan (available for download via Zenodo [45]). These old LD matrices were converted to the new CSR format using a custom script (see **Code Availability**), and the entries were cast and/or quantized from the standard double precision float (`float64`) to the four different data types listed in **Table** 1.

As for the GWAS summary statistics, we used the same summary data for Standing Height that was generated in our previous study [13]. In that analysis, we performed 5-fold cross validation on the White British subset in the UK Biobank (*N* = 337 225), where, for each fold, 80% of the samples were used for computing GWAS summary statistics and the remaining 20% were used as a heldout test set. Thus our main benchmarking results are shown as averages and standard errors across the 5 independent replicates of summary statistics for Standing Height in the UK Biobank [46].

#### 2.3.2 Benchmarking setup, metrics, and configurations

Our benchmarking experiments were focused on assessing the performance and resource usage of the `VIPRS` software along four axes. The first axis is the total wallclock time, which measures the time it takes to read the summary statistics files from disk, perform variant matching between the summary data and LD metadata, load the relevant entries of the LD matrix to memory, and perform variational inference to approximate the posterior for the effect sizes. Our optimizations should decrease the total wallclock time in three ways: (1) Reading the LD matrix from disk should be significantly faster, since stored entries are highly compressed. (2) The coordinate ascent updates should take considerably less time. (3) Parallelism, both within a coordinate ascent step and across chromosomes, should also reduce the total wallclock time. The second axis of performance is peak memory used throughout the program. This metric was measured using `\usr\bin\time` command on the Linux cluster where the experiments were done.

These first two measures are dependent on the characteristics of the data, convergence criteria, and the amount of computational resources used, which motivates the third axis: The runtime per iteration. This metric measures the time it takes to complete a single coordinate ascent step (also called the E-Step in our previous work [13]). Benchmarking the coordinate ascent step is done with a custom script (see **Code Availability**) where, after initializing the `VIPRS` module with data for a given chromosome, we invoke the coordinate ascent function *K* times to reach a total runtime of roughly 1 second. From this, we estimate the runtime-per-iteration. To obtain robust confidence intervals, we repeat this operation a total of 15 times.

The fourth and final axis is prediction accuracy for Standing Height on the held-out test set. Prediction accuracy was measured using the pseudo R-squared metric as implemented in the `viprs` package [13]. The metric requires access to GWAS summary statistics from the held-out test set of each fold.

Users of the software can also assess some of these axes of performance on their own personal computing environments, by specifying the `–-output-profiler-metrics` flag to the CLI script `viprs_fit`.

### 2.4 Large-scale PRS analyses with Pan-UK Biobank GWAS data

#### 2.4.1 Selection criteria for phenotypes from the Pan-UKB GWAS resource

The Pan-UKB is a comprehensive study that aims to elucidate the genetics of thousands of measured phenotypes from the UK Biobank across all ancestry groups [18]. Genome-wide association (GWA) summary statistics are available for 7221 phenotypes across 6 continental ancestry groups, in addition to LDSC heritability estimates and other quality-control (QC) flags [18] (see **Data Availability**). For our analysis, we selected continuous phenotypes that passed all the QC criteria defined by the Pan-UKB study in at least three different ancestry groups. These QC criteria included sample size checks and restricting to phenotypes with bounded LDSC heritability estimates that are significantly greater than zero. Applying these filters results in 75 high quality and heritable phenotypes that span 7 different categories, from blood biochemistry to anthropometric measures. The phenotypes, their codes, categories, description, and heritability estimates are listed in **Supplementary Table** S1.

#### 2.4.2 Extracting UK Biobank genotype and phenotype data

If polygenic scores are to be useful in clinical settings, they must be applicable to individuals of different ancestries. Towards this goal, we expanded the scope of our analyses and the pre-computed LD reference panels that we provide to encompass six ancestry groups, previously defined by the Pan-UK Biobank project [18] (Return 2442 from the UK Biobank dataset [46]). Specifically, Return 2442 provides PCA coordinates and covariates data for 448 216 diverse samples in the UK Biobank, in addition to a column assigning each sample to one of the following six continental ancestry groups: `AFR` (African), `AMR` (Admixed American), `CSA` (Cental and South Asian), `EAS` (East Asian), `EUR` (European), and `MID` (Middle Eastern). Out of these samples, we selected individuals who have no close relatives in the dataset (based on the `related` column provided), which reduced the final sample size to *N* = 382 329 individuals. The breakdown of sample sizes by ancestry is shown in **Table** 2.

**Table 2:** Sample sizes of unrelated individuals across the six ancestry groups defined by the Pan-UK Biobank Project.

| Ancestry group | Code | Sample size |
| --- | --- | --- |
| African | AFR | 6255 |
| Admixed American | AMR | 987 |
| Central and South Asian | CSA | 8284 |
| East Asian | EAS | 2700 |
| European | EUR | 362 446 |
| Middle Eastern | MID | 1567 |
| All samples | ALL | 382 239 |

We followed a similar variant quality control protocol as the Pan-UKB study [18]. Specifically, we selected bi-allelic variants with INFO score ≥ 0.8 and minor allele count (MAC) of at least 20 across the entire cohort. The only additional filter that we applied is excluding variants with ambiguous strand. This resulted in a total of 23 332 104 variants that were used in our largest downstream analysis.

To generate smaller subsets of variants for more computationally tractable workflows, we applied two additional filters. In the first case, we applied a minor allele frequency filter of MAF ≥ 0.1%, which resulted in a dataset of 13 772 714 variants. In the second case, in addition to the MAF filter, we restricted to the expanded HapMap3+ set generated by the authors of the `LDPred2` software [12, 47], which resulted in a dataset of 1 431 634 variants.

With these filters at hand, we used `plink2 v2.00a6LM` [17] to extract and transform the UK Biobank genotype dosage data from the provided `bgen` files [46] to `plink1 BED` file format, with a hard-call threshold of 0.1. We also used `plink1.9 v1.90b4.6` to annotate the extracted `bim` files with the centiMorgan distance, using the HapMap3 genetic map [48].

To evaluate the predictive performance of the inferred PRS models, we also extracted phenotype measurements for the 75 phenotypes selected earlier (**Supplementary Table** S1). The phenotypes were extracted from the tabular data of the UK Biobank resource and for all participants. If multiple measurements for the same phenotype were available (due to repeat visits), we selected the initial measurement. For the purposes of evaluation, the phenotypes were kept on their original scale. The only quality control filter that we applied for each phenotype was removing outliers, defined as measurements exceeding 3 standard deviations from the mean.

#### 2.4.3 Computing LD matrices for all Pan-UKB Populations

With the extracted UK Biobank genotype data, we then used our software package `magenpy` to compute LD matrices for the six ancestry groups listed in **Table** 2. Within each ancestry group, we restricted to variants with minor allele count (MAC) of at least 20. To explore the spectral properties of LD matrices and their impact on numerical stability and inference quality, we computed LD matrices using three different settings.

First, we computed LD matrices using both windowed (i.e. banded) and block-diagonal masks. Windowed masks estimate LD between all pairs of variants that are at most 3 centiMorgan apart on the same chromosome. Block-diagonal masks compute LD between all variants within the same LD block, defined by the `LDetect` software [49] (see **Data Availability**). The `LDetect` authors only published block-diagonal masks for African, Asian, and European samples, broadly defined [49]. Following previous work in this space [10], we used the European block boundaries for EUR and MID ancestry groups, African boundaries for the AFR and AMR ancestry groups, and Asian boundaries for the EAS and CSA ancestries.

Second, we estimated LD matrices using two strategies for dealing with missing genotype calls: Mean imputation (MI) or simply discarding missing observations, as done in the LD estimators provided by `plink1.9 v1.90b4.6` [17]. Third, to examine the impact of quantization on compressibility and the spectral properties of LD matrices, we stored the data using both `int8` and `int16` quantization. In the main analyses of the paper, we mainly used block-diagonal LD matrices that are computed with Mean Imputation (MI) for missing values and stored using `int8` quantization.

Due to the relatively small sample sizes of the minority populations, we only computed their LD matrices using the HapMap3+ subset of variants. For the European samples, we computed three sets of LD matrices corresponding to each of the variant sets described above. The largest variant set (INFO > 0.8) yielded pairwise correlations between 17 745 575 variants in European samples, with the reduction mainly due to the threshold MAC > 20. The number of variants and storage size of each of the LD matrices is shown in Table 3. While the variant sets overlap with one another, due to the size of the larger matrices, we created separate copies for each variant set. The HapMap3+ LD matrices for the six ancestry groups are available for download via `GitHub` and `Zenodo` (see **Data Availability**).

**Table 3:** LD matrix storage size for each ancestry group in the Pan-UKB, containing pairwise correlations between well-imputed variants (INFO *>* 0.8). For European samples, we show LD matrix storage size across three different variant sets. LD matrices were estimated using block-diagonal masks from `LDetect` [49] and stored using `int8` and `int16` quantization.

| Ancestry group | Variant set | Number of variants | LD matrix storage size |  |
| --- | --- | --- | --- | --- |
|  |  |  | int16 | int8 |
| AFR | HapMap3+ | 1 259 175 | 0.59 GB | 0.23 GB |
| AMR | HapMap3+ | 1 286 943 | 0.63 GB | 0.26 GB |
| CSA | HapMap3+ | 1 379 991 | 1.12 GB | 0.36 GB |
| EAS | HapMap3+ | 1 146 910 | 0.79 GB | 0.27 GB |
| MID | HapMap3+ | 1 307 269 | 0.94 GB | 0.36 GB |
| EUR | HapMap3+ | 1 431 634 | 1.01 GB | 0.31 GB |
| EUR | MAF > 0.1% | 13 492 321 | 70.72 GB | 13.79 GB |
| EUR | MAC > 20 | 17 745 575 | 118.91 GB | 20.66 GB |

#### 2.4.4 Training VIPRS on Pan-UKB GWAS summary statistics

We split the Pan-UKB GWAS summary statistics files for the 75 continuous phenotypes per ancestry and merged them with variant and phenotype metadata provided in the Pan-UKB manifest (see **Data Availability**). Some phenotypes did not have GWAS summary statistics for some ancestry groups, due to limited sample sizes or other QC criteria.

After the summary statistics files were merged and post-processed, we fit `VIPRS v1.0.2` on all the available summary data, each of which was matched with the appropriate ancestry-specific LD reference panel described earlier. By default, `VIPRS` was run by excluding long-range LD regions (`exclude-lrld` flag). To reduce memory usage, inference was carried out with the triangular LD mode and with the `dequantize-on-the-fly` flag, which dequantizes the entries of the LD matrix only when needed. By default, the coordinate ascent inference step was done using 4 threads for the HapMap3+ reference panel and 8 threads were used for the 13m (MAF>0.001) and 18m (MAC>20) reference panels.

#### 2.4.5 Cross-ancestry evaluation of inferred PRS models in the UK Biobank

Our previous work established that `VIPRS` is competitive with state-of-the-art PRS methods in terms of prediction accuracy and that relative performance of PRS models within the training cohort tends to correlate with their relative accuracy in other ancestries [13]. In this study, we focused on cross-population transferability to assess prediction accuracy within the UK Biobank. Given the six ancestry groups defined by the Pan-UKB, we trained a PRS model on GWAS summary statistics from one reference population, and then evaluated its prediction accuracy on the remaining five populations. In addition to helping assess transferability, this procedure ensures that the test set is distinct from the training set. Also, unlike cross-biobank evaluation, this procedure is less prone to batch effects in genotyping technology or phenotypic measurements, since all of the data was collected and preprocessed in the same manner. One potential limitation with this procedure is that cross-population transferability could be affected by many factors, such as differences in allele frequency, genetic architecture, and SNP heritability across ancestries. For instance, rare variants may not be shared across different ancestries, thus the actual gains in using larger variant sets from the training population may be undermined when transferring to the other populations.

Cross-population evaluation was carried out by first performing linear scoring on the entire genotype matrix for all samples using `plink2 v2.00a6LM` [17]. With these scores at hand, we then split the samples by the Pan-UKB population labels and evaluated the prediction accuracy within each population separately. The evaluation metric used in all our analyses is incremental *R*^2^, defined as the proportion of phenotypic variance explained by the PRS. By “incremental” we mean that before computing the *R*^2^ metric, the phenotype is residualized against a set of covariates, including sex, age, and the first 20 Principal Components (PCs). These covariates, in addition to the population labels, are provided in “Return 2442” of the UK Biobank [18, 46].

#### 2.4.6 Extracting CARTaGENE genotype and phenotype data and cross-biobank evaluation

To ensure that the trends we observed within the UK Biobank are replicable, we extended our analysis to the CARTaGENE Biobank, a prospective cohort with genotype data on *N* = 29, 337 participants from Quebec, Canada [50]. The majority ancestry among CARTaGENE participants is French-Canadian, although individuals from other ancestries are also represented.

For our analysis, we extracted genotype and phenotype data for all participants in the CARTaGENE cohort. The imputed genotype calls are provided by the biobank in `PGEN` file format [17]. We used `plink2 v2.00a6LM` [17] to perform linear scoring and generate polygenic scores for the CARTaGENE participants. The scoring pipeline for CARTaGENE was slightly different from the UKB, since the former used genome build `hg38` for variant calling, whereas the standard UKB dataset used `hg19`. To translate the scoring files for the CARTaGENE dataset, we used mapping files published by the UKB-PPP consortium [51] (see **Data Availability**).

For the purposes of evaluation, we used PCA coordinates and ancestry clustering results generated in a previous study that examined patterns of genetic variation in CARTaGENE [52]. In our analysis, we used the top 10 PCs, age, and sex as covariates when computing incremental R-squared metrics. To restrict evaluation to European samples, we used UMAP+HDBSCAN clusters corresponding to individuals of European ancestry (*N* = 27 043) [52].

To extract corresponding phenotype data, we manually matched 32 out of the 75 continuous phenotypes that we analyzed in the Pan-UKB. The phenotypes were matched based on their definitions and distributions in both cohorts (**Supplementary Figure** S9). Similar to our Pan-UKB pipeline, we excluded outlier phenotypic measurements, primarily those exceeding 3 standard deviations from the mean of each phenotype. Some of the phenotypes were measured on different scales, discretized differently, or showed clear shift in distribution between the CARTaGENE and the UK Biobank. We expect these heterogeneities in phenotypic distribution to affect the portability of UKB-derived PRSs to the CARTaGENE cohort (see **Discussion**).

## 3 Results

### 3.1 Massive gains in speed and efficiency with new VIPRS data structures and inference algorithms

To assess the overall impact of the algorithmic improvements introduced in the latest release of the `VIPRS` software (`v0.1`), we systematically compared its performance to previously published version (`v0.0.4`) across four axes of computational and statistical performance: total wallclock time, runtime-per-iteration, peak memory utilization, and prediction accuracy (see **Methods**). In addition to these four metrics, we compared the storage requirements of the new CSR LD matrix format against previously published baselines. The benchmarking experiments were carried out using 5-fold GWAS summary statistics for Standing Height from the UK Biobank (see **Data Availability**) [13]. The dataset contains summary statistics for ≈ 1.1 HapMap3 variants [13].

**Figure** 2a shows dramatic reductions in storage requirements for the new CSR LD matrix format, up to 54-fold improvement in space efficiency when using the `int8` data type to represent the entries of the LD matrix. Due to the elimination of redundancy by storing only the upper triangular portion of the matrix, even the double precision data type consumes 2.5 times less storage space than the old matrix format (`VIPRS v0.0.4`). Combining this triangular format with quantization allowed us to reduce storage requirements by up to an order of magnitude (**Figure** 2a) relative to published LD matrix resources from commonly used PRS methods [10–12]. Although quantization is a lossy compression technique, the resolution for `int8`, the most compact data type, is still below 0.01, a value small enough to be generally thresholded by many statistical genetics methods [9, 53].

Our previous study established that `VIPRS v0.0.4` is one of the fastest model-based PRS methods published to-date, achieving an average wallclock time of roughly 15 minutes on ≈ 1.1 million HapMap3 variants [13]. Here, we show that, on the same HapMap3 subset, the new version of the software (`v0.1`) is over six times faster in terms of overall wallclock time (**Figure** 2b), consumes at least three times less memory (**Figure** 2c), meanwhile achieving comparable prediction accuracy on Standing Height in the UK Biobank (**Figure** 2d). On a dataset of this scale, our benchmarks show that `VIPRS v0.1` converges in less than 60 seconds and consumes a modest 800MB of RAM at its peak. **Figure** 2e largely explains this improved efficiency, as it demonstrates more than an order-of-magnitude improvement in runtime-per-iteration in the new version of the software. This scaling does not necessarily translate to total wallclock time because there are other fixed costs beyond the inference step itself, mainly loading the data from disk, harmonizing LD and summary statistics, as well as loading libraries and other dependencies.

In these experiments, we used both versions of the software with default settings. This means that no multi-threading or parallel processing is in effect and the LD mode for `v0.1` is triangular. To analyze the impact of the other algorithmic changes, we conducted more detailed ablation experiments, which we turn to next.

### 3.2 Further improvements in scalability with parallelism and compact LD data representation

Scaling PRS methods to tens of millions of variants requires faster coordinate ascent routines with minimal memory footprint. To understand how the two layers of parallelism that we introduced impact the performance of `VIPRS`, we examined their impact along two of the axes discussed earlier: Runtime-per-iteration and wallclock time. The benchmarking experiments in this section were done using the same 5-fold GWAS summary statistics for standing height in the UK Biobank (see **Methods, Data Availability**) [13].

**Figure** 3a shows that parallel coordinate ascent significantly improves the runtime-per-iteration of the algorithm in a way that is commensurate with the number of threads. Scaling is not linear with the number of threads, however, and does vary between chromosomes (inset in **Figure** 3a and **Supplementary Figure** S4). We hypothesize that this is in part due to the fact all threads write to a shared *q* vector (Algorithms 1 and 2), which can lead to synchronization overhead. Additionally, in some cases, multi-threading performance may be limited by memory bandwidth. We expect bigger benefits of multi-threading for larger chromosomes with relatively small LD blocks, which is consistent with the trends we observe in **Figure** 3a and **Supplementary Figure** S4.

**Figure 3:**
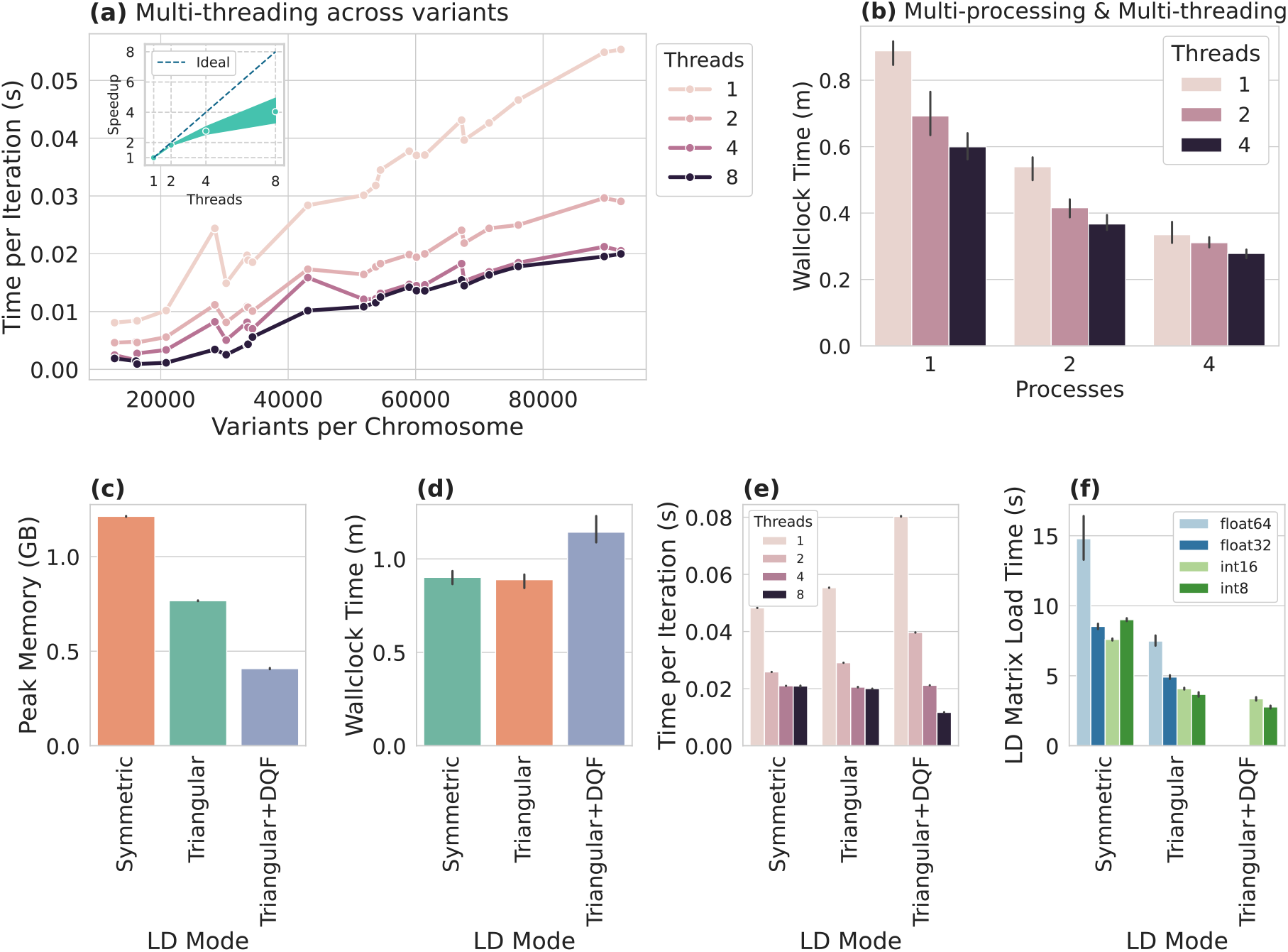
Illustrating the impact of multi-threading, parallel processing, and new Linkage-Disequilibrium (LD) matrix representations on the computational performance of `VIPRS`. Benchmarks are run on 5-fold GWAS summary statistics for Standing Height from the UK Biobank with ≈ 1.1 million HapMap3 variants. Panel **(a)** shows the runtime-per-iteration (in seconds) as a function of the number of variants on each chromosome (x-axis) and the number of threads used in the coordinate ascent updates (colors). The inset figure shows speedup in runtime-per-iteration as a function of the number of threads. Dots show average speedup and bands display smoothed 95% confidence interval across chromosomes. Panel **(b)** shows the total wallclock time (minutes) of the software as a function of the number of processes in the process pool (x-axis) and the number of threads used in the coordinate ascent step (colors). Panels **(c-d)** show the peak memory (GB) and total wallclock time (minutes) as a function of how the LD matrix is represented in memory (see **Methods**). Panel **(e)** shows and runtime-per-iteration (seconds) on Chromosome 1 as a function of the LD matrix representation (x-axis) and number of threads (colors). Panel **(f)** shows the time to load and post-process the LD matrix (seconds) for ≈ 1.1 million HapMap3 variants from disk to memory, across the three different modes (x-axis) and storage data types (colors). Vertical black lines above the bars in Panels **(b-d)** and **(f)** show standard error across the 5 folds.

The benefits of multi-threading on total wallclock time are evident, with four threads reducing total runtime by approximately 30% (**Figure** 3b). This improvement is achieved without affecting prediction accuracy on the held-out test set (**Supplementary Figure** S2). As discussed in the **Methods** section, the benefits of multi-threading on wallclock time may be partially dampened in some settings: multi-threaded implementations may require more iterations to reach convergence, especially when the variants/threads ratio is small. As for the second layer of parallelism, multi-processing across chromosomes with process pools shows clear benefits on wallclock time for this dataset, with the reductions in total runtime scaling proportionally with the number of processes (**Figure** 3b). This is to be expected, since processes have their own resources and copy of the relevant data and the main overhead here is forking the processes themselves.

The other set of algorithmic improvements introduced here are new memory-efficient implementations of the coordinate-ascent algorithm. **Figure** 3c shows that the Triangular LD mode outlined in Algorithm 2 significantly reduces memory usage compared to the symmetric LD mode (roughly 40% reduction), without affecting total wallclock time, runtime-per-iteration or prediction accuracy (**Figure** 3d-e, **Supplementary Figure** S2). Combining triangular LD mode with dequantizing the entries of the LD matrix on-the-fly (DQF) reduces memory utilization by another factor of two (**Figure** 3c), though the overhead of repeatedly dequantizing the data in every iteration results in notable slowdowns in both total wallclock time and runtime-per-iteration (**Figure** 3d-e). Interestingly, the DQF variant of the algorithm scales better with multi-threading in the coordinate ascent step (**Supplementary Figure** S4), perhaps due to reduced memory bandwidth requirement of the more compact LD data. Finally, **Figure** 3f shows the time to load the LD matrix depending on the encoded data type and LD mode for the algorithm. The benchmarks demonstrate that dequantizing the data in this case is very swift, though symmetrizing the matrix may approximately double the loading time.

Overall, in the case of a relatively small dataset of ≈ 1.1 million HapMap3 variants, the differences between these variations of the algorithm may seem small: Differences of 30 seconds are negligible and 600MB of RAM is insignificant for modern computing devices. Naturally though, we expect these relative differences to become consequential when performing inference over tens of millions of variants, as we discuss next.

### 3.3 Improved PRS transferability with inference over 18 million variants in the UK Biobank

To demonstrate the scalability of the new algorithmic improvements, we now turn to the Pan-UKB dataset, a comprehensive resource containing GWAS summary statistics for upwards of 7000 phenotypes across six continental ancestry groups [18, 46]. The quality control (QC) and association testing methodology is standardized across the entire dataset, making it an attractive resource for systematic PRS analyses. Furthermore, the Pan-UKB marginal association testing pipeline contains data for all well-imputed variants, up to 28 million, allowing us to test the computational performance and potential benefits of Bayesian modeling of the joint effects of more than a dozen million variants. In this section, we primarily focus on training `VIPRS` on GWAS data from European samples, as this group has the largest sample size in the UK Biobank.

#### 3.3.1 Computational efficiency and numerical stability

To assess the scalability of our method, we computed three new LD matrices corresponding to three sets of well-imputed variants in European samples (INFO > 0.8 [18]): 1.4 million HapMap3+ variants, 13.4 million variants (MAF > 0.001), and 18 million variants (MAC > 20) (**Table** 3, see **Methods**). We also extracted GWAS summary statistics for 75 of the most significantly heritable phenotypes in the UKB, based on quality control metrics provided by the Pan-UKB manifest (see **Supplementary Table** S1, **Data Availability**).

The first result to highlight is the tractability of the new compressed LD matrix format across the three variant sets. Combined with `int8` quantization, where entries are stored using a single byte, the proposed format results in highly compact LD matrices for the 1.4 million HapMap3+ variants, occupying less than 400MB of storage for each of the six ancestry groups defined in the Pan-UKB (**Table** 3, **Supplementary Table**S2). The advantages of this approach are even more apparent for denser variant sets, with the new format requiring only 13.8GB and 20.7GB of on-disk storage for matrices with up to 13.5 and 18 million variant set, respectively (**Table** 3). With `int16` quantization, LD matrices become less compressible, requiring up to 118.9GB of storage for the 18 million variants (**Table** 3). Nonetheless, this represents a vast improvement over the old storage format, which required >1 TiB of storage for half as many variants (9.8 million) [13]. It is also instructive to compare this approach with a recent method that decomposes LD matrices into a low-rank representation, which required up to 75 GB of storage for only 7 million variants [15]. This compressibility should make it easier to store and disseminate large-scale LD matrices, at low cost and with little extra overhead.

With these highly compressed LD matrices at hand, we fit the new `VIPRS v0.1` method to the Pan-UKB GWAS summary statistics for the 75 phenotypes described earlier across the three variant sets in European samples. In this analysis, we first examined computational performance as well as the numerical stability and convergence of the inference algorithm. Since LD data storage type (`int8` vs. `int16`) had negligible impact on prediction accuracy (**Supplementary Figure** S13), we mainly present results from analyses using the `int8` data type, which is more computationally tractable (**Supplementary Figure** S6). **Figure** 4a shows that `VIPRS v0.1` converges in less than an hour on GWAS summary data for up to 18 million variants. With the dequantize-on-the-fly (DQF) option introduced in the latest software implementation, the method requires less than 15GB of memory at its peak for these datasets (**Figure** 4b). We highlight that, in these experiments, the runtime of the program is affected by fetching the LD data across the cluster network (teal color **Figure** 4a), a task that should have negligible runtime if the data is stored locally on the compute node. The task of training/inference (orange color in **Figure** 4a) requires roughly 40 minutes genome-wide. Additionally, these results are obtained while utilizing modest computational resources: 8 CPU cores. When run on High-performance computing (HPC) clusters with over 40 cores, the algorithm converges in less than 20 minutes genome-wide (**Supplementary Figure** S5).

**Figure 4:**
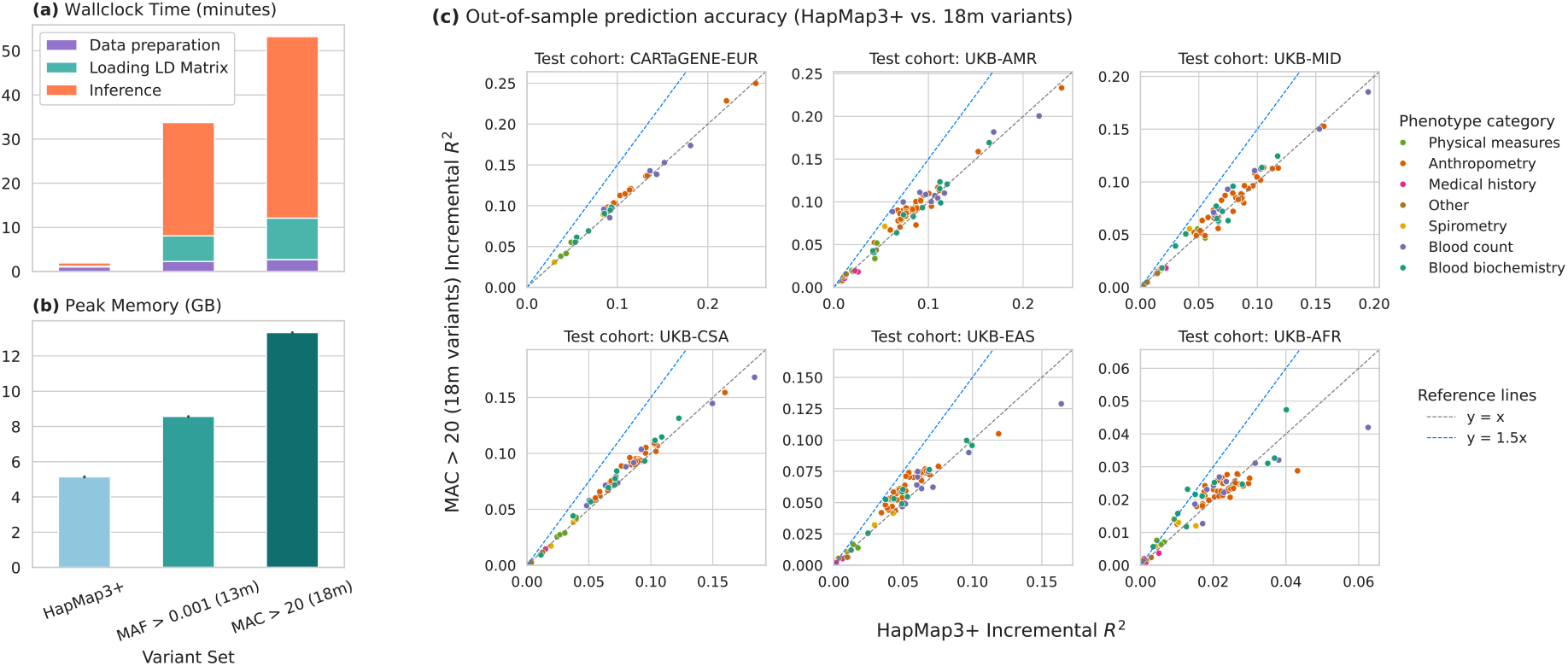
Systematic evaluation of PanUKB-derived PRS models across 75 continuous phenotypes and three variant sets. All polygenic scores were inferred from European GWAS summary statistics from the PanUKB initiative using `VIPRS v0.1`. Panels **(a-b)** show the average wallclock time (minutes) and peak memory usage (GB) across all the phenotypes. Colors in panel **(a)** denote average time required for each sub-task during inference. Panel **(c)** shows the comparative prediction accuracy (Incremental R-squared) between the HapMap3+ variant set on the x-axis (1.4 million variants) compared to an expanded variant set including up to 18 million variants on the y-axis. Each sub-panel compares the accuracy for one of six held-out test cohorts across two biobanks: CARTaGENE and UK Biobank (UKB). The ancestry groups are EUR (European), AMR (Admixed American), MID (Middle Eastern), CSA (Central and South Asian), EAS (East Asian), and AFR (African). Colors in panel **(c)** denote different phenotype categories and dashed lines delineate the magnitude of the improvement in prediction accuracy.

Given the potential for numerical instabilities discussed in **Methods**, we examined model convergence when performing inference in ultra high dimensions. In **Supplementary Figure** S10, we show that the incremental R-squared metric on the training set improves uniformly across the 75 phenotypes examined. This supports the idea that the algorithm is stable and the model converges well in these high dimensional settings, though it does not necessarily imply that these larger models will extrapolate better.

#### 3.3.2 Accuracy and transferability of highly parametrized PRS models

To test the extrapolation ability of the inferred PRS models with the three variant sets, we assessed the prediction accuracy on six held-out test cohorts across two biobanks (see **Methods**). In the first case, we examined within-ancestry, but cross-biobank transferability, where UKB-derived PRS models trained on European GWAS data were evaluated on European samples in the CARTaGENE biobank [50]. In the second case, we examined the transferability of those same UKB-EUR PRS models on the five held-out ancestry groups in the Pan-UKB (**Table** 2) [18].

In the cross-biobank analysis, we observed modest improvements in prediction accuracy when using denser variant sets, with an average improvement of 3% across the 32 phenotypes that were matched between the two biobanks (**Figure** 4c and **Supplementary Figure** S11;S12). The marginal benefit provided by the denser variant sets in this setting suggests that the HapMap3+ variant set tags most causal variants in Europeans quite well. It might also be the case that these observations may be indicative of slight overfitting, though the scale of improvement is consistent with recent analyses [15] (see **Discussion**).

However, in the cross-ancestry case, it appears that denser variant sets provide clear benefit in portability, with up to 50% improvement in prediction accuracy for many phenotypes and ancestry groups (**Figure** 4c and **Supplementary Figure** S11;S12). The scale of improvement, 10-15% on average across the five ancestry groups, is on-par with what is sometimes reported by cross-ancestry PRS methods, such as `PRS-CSx` [54] and is consistent with recent analyses examining the potential benefits of modeling dense variant sets [15]. This improved portability, when contrasted with the cross-biobank performance, also highlights the importance of capturing the true causal variants for accurate cross-ancestry PRS analyses.

## 4 Discussion

In this work, we presented an integrated suite of data structures, algorithms, and software tools to scale model-based PRS inference to millions of genetic variants, meeting the ever-increasing depth and resolution of modern GWAS studies [55]. Our technical contributions extend from the practical solutions for large-scale LD matrix computation and storage to robust and efficient coordinate-wise optimization algorithms that can perform Variational Bayesian regression [13] over a dozen million variants in as little as 20 minutes and using less than 15 Gigabytes of RAM.

LD matrices are an integral part of modern statistical genetics toolkits, playing an essential role in fine-mapping [24,56], PRS inference [8,10–13,15,27], and imputation or simulation of summary statistics [19,57], among other applications. Previous work has explored various techniques for computing, sparsifying, and transforming LD matrices [8,11, 13,16, 23,27, 58], with the aims of making downstream inference algorithms more efficient, numerically stable, and accurate. Despite this, clear and universal standards for representing LD matrices are lacking, leading each new method to develop its own set of tools and storage formats. In this work, we attempted to pool together best practices from disparate subfields of statistical genetics to present a universal, cloud-native storage format [26], with compression ratios of up to 50 fold over naive approaches. This compactness rivals recent low-rank representations of LD matrices [15] while being a more universal representation that can readily serve a wide variety of applications. It is also competitive with a recently-proposed, ultra sparse estimator of precision matrices (inverse of LD) [19], making it useful for many applications that require access to pairwise correlations between genetic variants. Compressibility of LD matrices provides many practical benefits besides allowing for more efficient inference algorithms. For one, it reduces the barrier for computing and releasing LD matrices for fine-grained subgroups in large cohorts or biobanks, potentially improving the quality of PRS inference in some minority populations [16]. Recent efforts to release comprehensive LD matrices as a public resource proved to be costly, requiring Terabytes of storage [18, 59]. While cloud storage costs are relatively minor, moving data of this scale along global or local networks can still incur substantial costs.

Furthermore, our analysis highlighted the importance of the spectral properties of LD matrices on the accuracy and numerical stability of PRS inference algorithms [40, 41]. Concretely, our analyses outlined how the spectral properties of an LD matrix are affected by many common choices for estimating pairwise correlations between genetic variants. These choices include the definition of sparsification mask (e.g. banded or block-diagonal), estimation in the presence of missing genotype calls, inclusion of variants in long-range LD regions, thresholding, and approximation errors. While we provided practical recommendations for computing well-conditioned matrices that behave stably even in ultra high-dimensions, there remains many open questions with regards to the precise balance between compression, shrinkage, and accuracy that we leave for future work.

The compressed LD matrix format integrates seamlessly with the new coordinate ascent inference algorithms, allowing us to derive efficient coordinate-wise update rules that operate on compressed and non-redundant LD data. This “low-memory” version of the new `VIPRS` method reduces memory usage during inference by more than order of magnitude compared to a naive implementation while achieving comparable prediction accuracy. Our work further demonstrates that these update rules can be easily parallelized without assuming any special structure, such as independent LD blocks, leading to significant improvements in inference speed. Notably, we found that combining parallel coordinate ascent with compact LD representations, i.e. quantization, not only saved memory but also enhanced scaling performance (**Figure** 3; **Supplementary Figure** S4).

This tangible boost in processing speed and memory efficiency enabled testing the limits of model-based PRS inference in a systematic manner, using GWAS summary statistics for up to 18 million well-imputed variants across 75 of the most heritable phenotypes in the Pan-UKB [18, 46]. Cross-population validation analyses within the UKB demonstrated that going beyond the standard HapMap3 subset robustly improves prediction accuracy across most populations and phenotype categories. Due to the Bayesian framework of our method, these improvements are especially notable in light of the overfitting observed in a recent study examining the effects of applying more permissive thresholds in the heuristic Clumping-and-Thresholding (C+T) approach for polygenic score construction [60].

We hypothesize that these gains in cross-population prediction accuracy are mainly due to one of two factors. First, expanding the variant set makes it more likely to incorporate causal variants not well-tagged by the HapMap3 subset. This includes relatively rare variants with large effect on the phenotype of interest. Second, given differences in LD structure across populations [61], even if true causal variants are well-tagged by the HapMap3 subset in European samples, a model that incorporates true causal variants in the regression is likely to transfer better than a model using tagging variants alone. While the cross-population validation analyses yielded clear benefits in transferability, the scale of the improvement within the same ancestry group seemed modest. This suggests that, in European samples at least, the HapMap3+ variants tag most causal variants for these phenotypes rather well. Other factors could have dampened the impact of using more dense variant sets. For instance, many of the manually-matched phenotypes showed significant differences in their distribution across the two biobanks (**Supplementary Figure** S9), introducing potential batch effects. Furthermore, while both biobanks used similar genotyping arrays for their respective cohorts [46, 50], imputation pipelines and reference panels are different [46, 50], which might lead to systematic biases in the imputation of some causal variants. Nonetheless, `VIPRS` still demonstrates overall robust performance in the cross-biobank setting and the scale of the improvement we report here is consistent with previous studies [15].

Finally, many of the gains in efficiency and scalability presented here are not restricted to the `VIPRS` inference algorithms per se and should be widely applicable. Many coordinate-wise optimization or sampling-based methods, such as `Lassosum` [8] and `LDPred` [12, 27], can benefit from the “low-memory” updating scheme in addition to unrestricted parallelism to improve computational efficiency. Although parallel coordinate descent algorithms for the Lasso have been extensively studied [34], there is also comparable, though less well known, literature on parallel Gibbs sampling algorithms [62]. The cloud-native `Zarr` format [26] which powers our LD matrix storage specification, comes with public APIs in many popular programming languages, which should make it relatively easy to access and build on the LD storage format presented in this work.

In summary, our new `VIPRS` software offers orders of magnitude improvements in storage, computation, and memory efficiency, requiring only 20 minutes to infer the joint effects of all well-imputed bi-allelic variants in the UK Biobank. These improvements situate it as the next-generation PRS tool in the modern pipeline of increasingly large-scale and comprehensively phenotyped and genotyped biobanks.

## Code Availability

- Scripts to replicate all the analyses and generate the figures for this manuscript are available via GitHub: https://github.com/shz9/viprs-benchmarks-paper
- The `magenpy` python package for computing LD matrices / harmonizing LD and GWAS summary statistics: https://github.com/shz9/magenpy
- Latest version of the `VIPRS` software: https://github.com/shz9/viprs
- A Google Colab notebook illustrating the speed and main features of the commandline interface (CLI) of the `viprs` software: https://github.com/shz9/viprs/blob/master/notebooks/viprs_cli_example.ipynb
- Script to convert between old and new format for the `magenpy` /`VIPRS` LD matrices: https://github.com/shz9/magenpy/blob/master/examples/convert_old_ld_matrices.py

## Data Availability

- LD matrices for the six continental ancestry groups are available for download via GitHub (https://github.com/shz9/viprs/releases/tag/v0.1.2) and Zenodo (https://zenodo.org/records/14614207).
- Five-fold benchmarking summary statistics for Standing Height: https://doi.org/10.5281/zenodo.14270953
- LD blocks defined by `LDetect` : https://bitbucket.org/nygcresearch/ldetect-data/src/master/
- The Pan-UKB phenotype manifest with heritability estimates, QC flags, and hyperlinks to download GWAS summary statistics: https://docs.google.com/spreadsheets/d/1AeeADtT0U1AukliiNyiVzVRdLYPkTbr
- Mapping files for rsids across genome builds `hg19` and `hg38` from the UKB-PPP: https://www.synapse.org/Synapse:syn51364943/wiki/622119

## Acknowledgements

We thank members of the Li and Gravel labs for useful feedback and discussions on earlier drafts of this manuscript. Y.L. is supported by Canada Research Chair (Tier 2) in Machine Learning for Genomics and Healthcare (CRC-2021-00547) and Natural Sciences and Engineering Research Council (NSERC) Discovery Grant (RGPIN-2016-05174). This research was supported by the Canadian Institute for Health Research (CIHR) project grant 437576, NSERC grant RGPIN-2017-04816, the Canada Research Chair program to S.G., and the Canada Foundation for Innovation. This research used the NeuroHub infrastructure and was undertaken thanks in part to funding from the Canada First Research Excellence Fund, awarded through the Healthy Brains, Healthy Lives initiative at McGill University. This research was enabled in part by support provided by Calcul Québec and the Digital Research Alliance of Canada. This research has been conducted using the UK Biobank Resource under Application Number 45551.

## Author contributions

Y.L., S.G. and S.M. supervised the work. Under the supervision of Y.L. and S.M., S.Z. and C.A.H benchmarked the software, analyzed computational bottlenecks, and tested various solutions to speed up the coordinate ascent step. S.Z. implemented the new LD storage format and associated data structures and software utilities. S.G. and S.Z. examined the spectral properties of LD matrices and their impact on model convergence and stability. C.A.H. and S.Z. conducted the benchmarking experiments. S.Z. conducted the Pan-UKB analyses. S.Z. wrote the initial manuscript. All authors edited and reviewed the final version.

## Declaration of Interests

The authors declare no competing interests.

## Notes

https://github.com/shz9/viprs-benchmarks-paper

### Competing Interest Statement

The authors have declared no competing interest.

## References

[1] Natarajan, P. et al. Polygenic risk score identifies subgroup with higher burden of atherosclerosis and greater relative benefit from statin therapy in the primary prevention setting. Circulation 135 (2017).

[2] Khera, A. V. et al. Genome-wide polygenic scores for common diseases identify individuals with risk equivalent to monogenic mutations. Nature Genetics 50 (2018).

[3] Sugrue, L. P. & Desikan, R. S. What are polygenic scores and why are they important? JAMA - Journal of the American Medical Association 321 (2019).

[4] Lewis, C. M. & Vassos, E. Polygenic risk scores: from research tools to clinical instruments. Genome Medicine 12, 44 (2020). URL 10.1186/s13073-020-00742-5.

[5] Pain, O. et al. Evaluation of polygenic prediction methodology within a reference-standardized framework. PLOS Genetics 17, 1–22 (2021). URL 10.1371/journal.pgen.1009021.

[6] Yang, S. & Zhou, X. PGS-server: accuracy, robustness and transferability of polygenic score methods for biobank scale studies. Briefings in Bioinformatics 23 (2022). URL 10.1093/bib/bbac039.Bbac039, https://academic.oup.com/bib/article-pdf/23/2/bbac039/42806392/bbac039.pdf.

[7] Jayasinghe, D., Eshetie, S., Beckmann, K., Benyamin, B. & Lee, S. H. Advancements and limitations in polygenic risk score methods for genomic prediction: a scoping review. Human Genetics (2024). URL 10.1007/s00439-024-02716-8.

[8] Mak, T. S. H., Porsch, R. M., Choi, S. W., Zhou, X. & Sham, P. C. Polygenic scores via penalized regression on summary statistics. Genetic Epidemiology 41 (2017).

[9] Choi, S. W. & O’Reilly, P. F. Prsice-2: Polygenic risk score software for biobank-scale data. GigaScience 8 (2019).

[10] Ge, T., Chen, C. Y., Ni, Y., Feng, Y. C. A. & Smoller, J. W. Polygenic prediction via bayesian regression and continuous shrinkage priors. Nature Communications 10 (2019).

[11] Lloyd-Jones, L. R. et al. Improved polygenic prediction by bayesian multiple regression on summary statistics. Nature Communications 10 (2019).

[12] Privé, F., Arbel, J. & Vilhjálmsson, B. J. Ldpred2: Better, faster, stronger. Bioinformatics 36 (2020).

[13] Zabad, S., Gravel, S. & Li, Y. Fast and accurate bayesian polygenic risk modeling with variational inference. The American Journal of Human Genetics 110, 741–761 (2023). URL https://www.sciencedirect.com/science/article/pii/S0002929723000939.

[14] Lambert, S. A. et al. The polygenic score catalog as an open database for reproducibility and systematic evaluation. Nature Genetics 53, 420–425 (2021). URL 10.1038/s41588-021-00783-5.

[15] Zheng, Z. et al. Leveraging functional genomic annotations and genome coverage to improve polygenic prediction of complex traits within and between ancestries. Nature Genetics 56, 767–777 (2024). URL 10.1038/s41588-024-01704-y.

[16] Privé, F., Arbel, J., Aschard, H. & Vilhjálmsson, B. J. Identifying and correcting for misspecifications in gwas summary statistics and polygenic scores. Human Genetics and Genomics Advances 3 (2022). URL 10.1016/j.xhgg.2022.100136.

[17] Chang, C. C. et al. Second-generation plink: Rising to the challenge of larger and richer datasets. GigaScience 4 (2015).

[18] Karczewski, K. J. et al. Pan-uk biobank gwas improves discovery, analysis of genetic architecture, and resolution into ancestry-enriched effects. medRxiv (2024). URL https://www.medrxiv.org/content/early/2024/03/15/2024.03.13.24303864. https://www.medrxiv.org/content/early/2024/03/15/2024.03.13.24303864.full.pdf.

[19] Salehi Nowbandegani, P. et al. Extremely sparse models of linkage disequilibrium in ancestrally diverse association studies. Nature Genetics 55, 1494–1502 (2023). URL 10.1038/s41588-023-01487-8.

[20] Zhang, Q., Privé, F., Vilhjálmsson, B. & Speed, D. Improved genetic prediction of complex traits from individual-level data or summary statistics. Nature Communications 12, 4192 (2021). URL 10.1038/s41467-021-24485-y.

[21] Hail Team, H. T. Hail. URL https://github.com/hail-is/hail.

[22] Benner, C. et al. Prospects of fine-mapping trait-associated genomic regions by using summary statistics from genome-wide association studies. The American Journal of Human Genetics 101, 539–551 (2017). URL https://www.sciencedirect.com/science/article/pii/S0002929717303348.

[23] Weiner, R. J., Lakhani, C., Knowles, D. A. & Gürsoy, G. LDmat: efficiently queryable compression of linkage disequilibrium matrices. Bioinformatics 39, btad092 (2023). URL 10.1093/bioinformatics/btad092. https://academic.oup.com/bioinformatics/article-pdf/39/2/btad092/49345179/btad092.pdf.

[24] Benner, C. et al. FINEMAP: efficient variable selection using summary data from genome-wide association studies. Bioinformatics 32, 1493–1501 (2016). URL 10.1093/bioinformatics/btw018. https://academic.oup.com/bioinformatics/article-pdf/32/10/1493/49019673/bioinformatics_32_10_1493.pdf.

[25] Wu, H., Judd, P., Zhang, X., Isaev, M. & Micikevicius, P. Integer quantization for deep learning inference: Principles and empirical evaluation (2020). 2004.09602.

[26] Zarr python. https://zarr.readthedocs.io/en/stable/. Accessed: 2024-05-29.

[27] Vilhjálmsson, B. J. et al. Modeling linkage disequilibrium increases accuracy of polygenic risk scores. American Journal of Human Genetics 97 (2015).

[28] Blackford, L. S. et al. An updated set of basic linear algebra subprograms (blas). ACM Transactions on Mathematical Software 28, 135–151 (2002).

[29] Yang, S. & Zhou, X. Accurate and scalable construction of polygenic scores in large biobank data sets. The American Journal of Human Genetics 106, 679–693 (2020). URL https://www.sciencedirect.com/science/article/pii/S0002929720301099.

[30] Joblib: running python functions as pipeline jobs. https://joblib.readthedocs.io/en/stable/. Accessed: 2024-05-29.

[31] OpenMP Architecture Review Board. OpenMP application program interface version 3.0 (2008). URL http://www.openmp.org/mp-documents/spec30.pdf.

[32] Richtárik, P. & Takáč, M. Parallel coordinate descent methods for big data optimization. Mathematical Programming 156, 433–484 (2016). URL 10.1007/s10107-015-0901-6.

[33] Plummer, S., Pati, D. & Bhattacharya, A. Dynamics of coordinate ascent variational inference: A case study in 2d ising models. Entropy 22 (2020). URL https://www.mdpi.com/1099-4300/22/11/1263.

[34] Liu, J., Wright, S., Re, C., Bittorf, V. & Sridhar, S. An asynchronous parallel stochastic coordinate descent algorithm. In Xing, E. P. & Jebara, T. (eds.) Proceedings of the 31st International Conference on Machine Learning, vol. 32 of Proceedings of Machine Learning Research, 469–477 (PMLR, Bejing, China, 2014). URL https://proceedings.mlr.press/v32/liud14.html.

[35] Merkel, D. Docker: lightweight linux containers for consistent development and deployment. Linux journal 2014, 2 (2014).

[36] Chen, W. et al. Improved analyses of gwas summary statistics by reducing data heterogeneity and errors. Nature Communications 12, 7117 (2021). URL 10.1038/s41467-021-27438-7.

[37] Virtanen, P. et al. SciPy 1.0: Fundamental Algorithms for Scientific Computing in Python. Nature Methods 17, 261–272 (2020).

[38] Lehoucq, R. B., Sorensen, D. C. & Yang, C. ARPACK: Solution of Large Scale Eigenvalue Problems by Implicitly Restarted Arnoldi Methods. Available from (1997).

[39] George, R. K. & Ajayakumar, A. A Course in Linear Algebra, vol. 35 of University Texts in the Mathematical Sciences (2024).

[40] Choi, Y. & Tibshirani, R. An investigation of methods for handling missing data with penalized regression (2013). URL https://arxiv.org/abs/1310.2076.1310.2076.

[41] Loh, P.-L. & Wainwright, M. J. High-dimensional regression with noisy and missing data: Provable guarantees with nonconvexity. The Annals of Statistics 40, 1637–1664 (2012). URL 10.1214/12-AOS1018.

[42] Datta, A. & Zou, H. CoCoLasso for high-dimensional error-in-variables regression. The Annals of Statistics 45, 2400–2426 (2017). URL 10.1214/16-AOS1527.

[43] Escribe, C. et al. Block coordinate descent algorithm improves variable selection and estimation in error-in-variables regression. Genetic Epidemiology 45, 874–890 (2021). URL https://onlinelibrary.wiley.com/doi/abs/10.1002/gepi.22430. https://onlinelibrary.wiley.com/doi/pdf/10.1002/gepi.22430.

[44] Spence, J. P., Sinnott-Armstrong, N., Assimes, T. L. & Pritchard, J. K. A flexible modeling and inference framework for estimating variant effect sizes from gwas summary statistics. bioRxiv (2022). URL https://www.biorxiv.org/content/early/2022/04/19/2022.04.18.488696. https://www.biorxiv.org/content/early/2022/04/19/2022.04.18.488696.full.pdf.

[45] Zabad, S., Gravel, S. & Li, Y. LD matrices from the White British cohort in the UK Biobank in Zarr format (2022). URL 10.5281/zenodo.7036625.

[46] Bycroft, C. et al. The uk biobank resource with deep phenotyping and genomic data. Nature 562 (2018).

[47] Privé, F., Albiñana, C., Arbel, J., Pasaniuc, B. & Vilhjálmsson, B. J. Inferring disease architecture and predictive ability with ldpred2-auto. The American Journal of Human Genetics 110, 2042–2055 (2023). URL https://www.sciencedirect.com/science/article/pii/S0002929723003634.

[48] Altshuler, D. M. et al. Integrating common and rare genetic variation in diverse human populations. Nature 467 (2010).

[49] Berisa, T. & Pickrell, J. K. Approximately independent linkage disequilibrium blocks in human populations. Bioinformatics 32 (2016).

[50] Awadalla, P. et al. Cohort profile of the CARTaGENE study: Quebec’s population-based biobank for public health and personalized genomics. International Journal of Epidemiology 42, 1285–1299 (2012). URL 10.1093/ije/dys160. https://academic.oup.com/ije/article-pdf/42/5/1285/18481769/dys160.pdf.

[51] Sun, B. B. et al. Plasma proteomic associations with genetics and health in the uk biobank. Nature 622, 329–338 (2023). URL 10.1038/s41586-023-06592-6.

[52] Diaz-Papkovich, A. et al. Topological stratification of continuous genetic variation in large biobanks. bioRxiv (2023). URL https://www.biorxiv.org/content/early/2023/07/07/2023.07.06.548007. https://www.biorxiv.org/content/early/2023/07/07/2023.07.06.548007.full.pdf.

[53] Privé, F., Vilhjálmsson, B. J., Aschard, H. & Blum, M. G. Making the most of clumping and thresholding for polygenic scores. The American Journal of Human Genetics 105, 1213–1221 (2019). URL https://www.sciencedirect.com/science/article/pii/S0002929719304227.

[54] Ruan, Y. et al. Improving polygenic prediction in ancestrally diverse populations. Nature Genetics 54, 573–580 (2022). URL 10.1038/s41588-022-01054-7.

[55] Li, S., Carss, K. J., Halldorsson, B. V. & Cortes, A. Whole-genome sequencing of half-a-million uk biobank participants. medRxiv (2023). URL https://www.medrxiv.org/content/early/2023/12/08/2023.12.06.23299426. https://www.medrxiv.org/content/early/2023/12/08/2023.12.06.23299426.full.pdf.

[56] Zou, Y., Carbonetto, P., Wang, G. & Stephens, M. Fine-mapping from summary data with the “sum of single effects” model. PLOS Genetics 18, 1–24 (2022). URL 10.1371/journal.pgen.1010299.

[57] Pasaniuc, B. et al. Fast and accurate imputation of summary statistics enhances evidence of functional enrichment. Bioinformatics 30, 2906–2914 (2014). URL 10.1093/bioinformatics/btu416. https://academic.oup.com/bioinformatics/article-pdf/30/20/2906/48929857/bioinformatics_30_20_2906.pdf.

[58] Wen, X. & Stephens, M. Using linear predictors to impute allele frequencies from summary or pooled genotype data. Annals of Applied Statistics 4 (2010).

[59] Weissbrod, O. et al. Leveraging fine-mapping and multipopulation training data to improve cross-population polygenic risk scores. Nature Genetics 54, 450–458 (2022). URL 10.1038/s41588-022-01036-9.

[60] Aw, A. J., McRae, J., Rahmani, E. & Song, Y. S. Highly parameterized polygenic scores tend to overfit to population stratification via random effects. bioRxiv (2024). URL https://www.biorxiv.org/content/early/2024/01/29/2024.01.27.577589. https://www.biorxiv.org/content/early/2024/01/29/2024.01.27.577589.full.pdf.

[61] Evans, D. M. & Cardon, L. R. A comparison of linkage disequilibrium patterns and estimated population recombination rates across multiple populations. The American Journal of Human Genetics 76, 681–687 (2005). URL https://www.sciencedirect.com/science/article/pii/S0002929707628791.

[62] Johnson, M. J., Saunderson, J. & Willsky, A. Analyzing hogwild parallel gaussian gibbs sampling. In Burges, C., Bottou, L., Welling, M., Ghahramani, Z. & Weinberger, K. (eds.) Advances in Neural Information Processing Systems, vol. 26 (Curran Associates, Inc., 2013). URL https://proceedings.neurips.cc/paper_files/paper/2013/file/b51a15f382ac914391a58850ab343b00-Paper.pdf.

