## Supplementary for "Towards whole-genome inference of polygenic scores with fast and memory-efficient algorithms"

### S1 Supplementary Figures

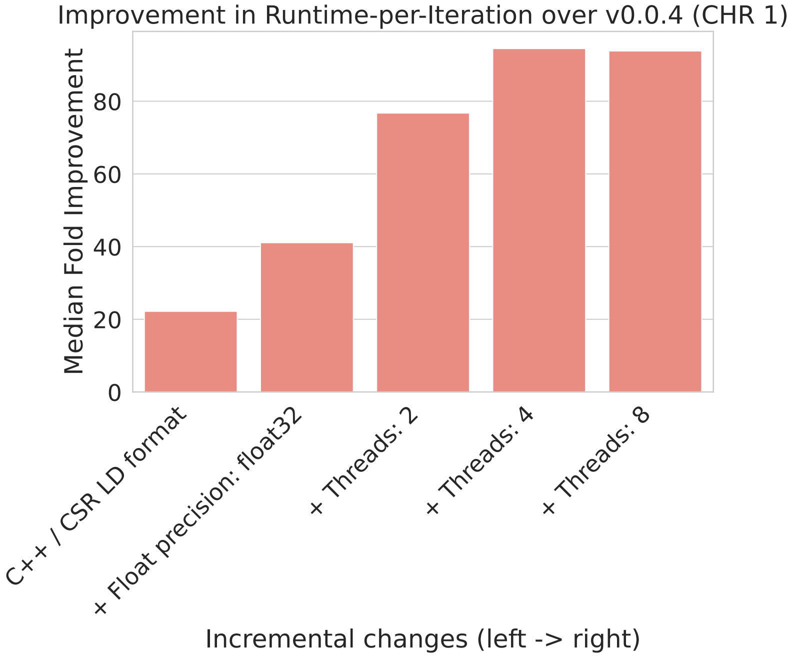


Figure S1: Median relative improvement in the runtime-per-iteration for Chromosome 1 in `VIPRS` v0.1 over `VIPRS` v0.0.4 (y-axis) as a function of the incremental updates to the software, from left to right (x-axis). The figure summarizes the impact of the various optimizations when chained together, starting from the new layout for the LD matrix / new implementation of E-Step in `C++`, proceeding to using single precision floating point format for the parameters, and ends with gains due to the multi-threaded implementation (2, 4, and 8 threads). Median improvement is computed by taking the median runtime of `VIPRS` v0.0.4 across 15 replicates and dividing by the median runtime of `VIPRS` v0.1 across the same number of replicates.

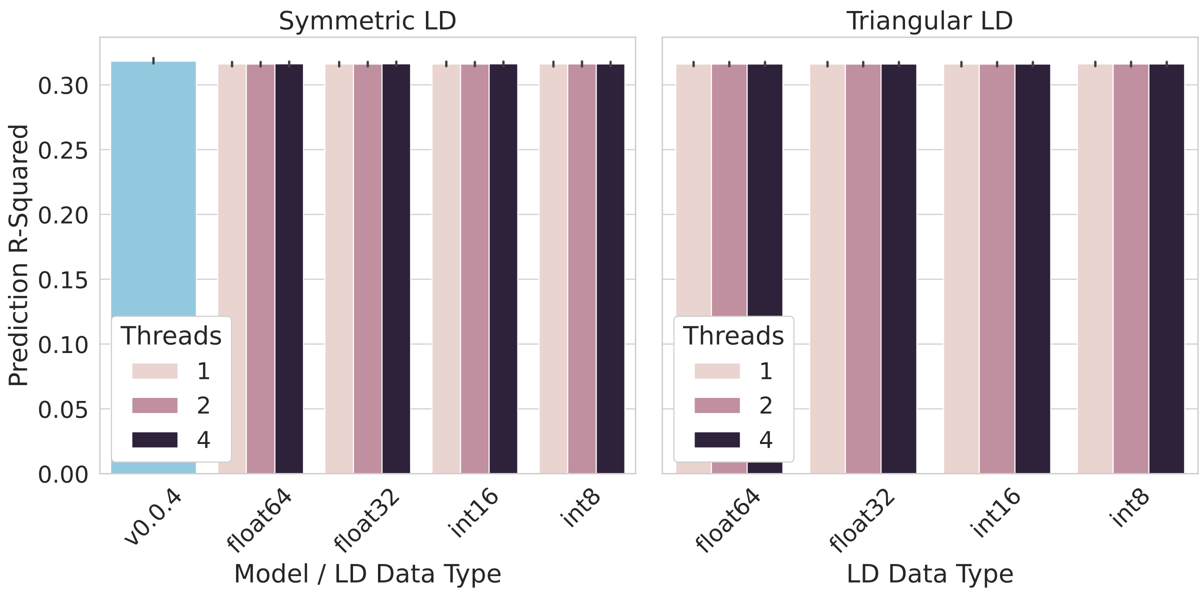


Figure S2: Prediction accuracy for Standing Height in the UK Biobank as a function of LD Mode / LD Data Type / and the number of threads used in the Coordinate Ascent step of `VIPRS` v0.1. The left panel shows Prediction R-Squared for the symmetric LD version of the algorithm and the right panels shows the same metric for the triangular LD version of the algorithm. X-axis shows the LD data types (`float64`, `float32`, `int16`, `int8`) as well as the v0.0.4 version of the software on the far left of the left panel. Threads used in the multi-threaded implementation of `VIPRS` v0.1 are shown in color. Black lines above the bars show standard error across the 5 folds.

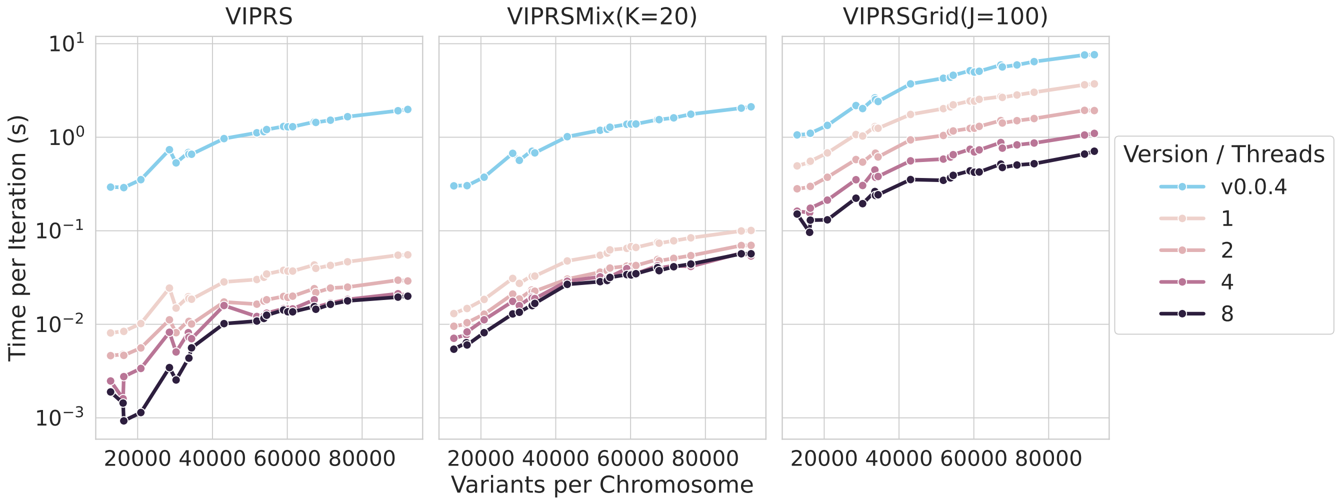


Figure S3: Mean runtime-per-iteration (in second and on log-scale) as a function of the number of SNPs for all the three configurations of the `VIPRS` model. The lines show the performance across versions v0.0.4 (skyblue) and v0.1 with different number of threads (color gradient). Each panel shows a different setup/configuration of the `VIPRS` model. Leftmost panel shows the runtime-per-iteration of the vanilla `VIPRS` with the spike-and-slab prior. Second panel in the middle shows the runtime-per-iteration of `VIPRS(K=20)`, which uses the sparse mixture prior with 20 components. Rightmost panel shows the runtime-per-iteration of the `VIPRSGrid(J=100)` model, which performs inference over 100 different hyperparameter settings.

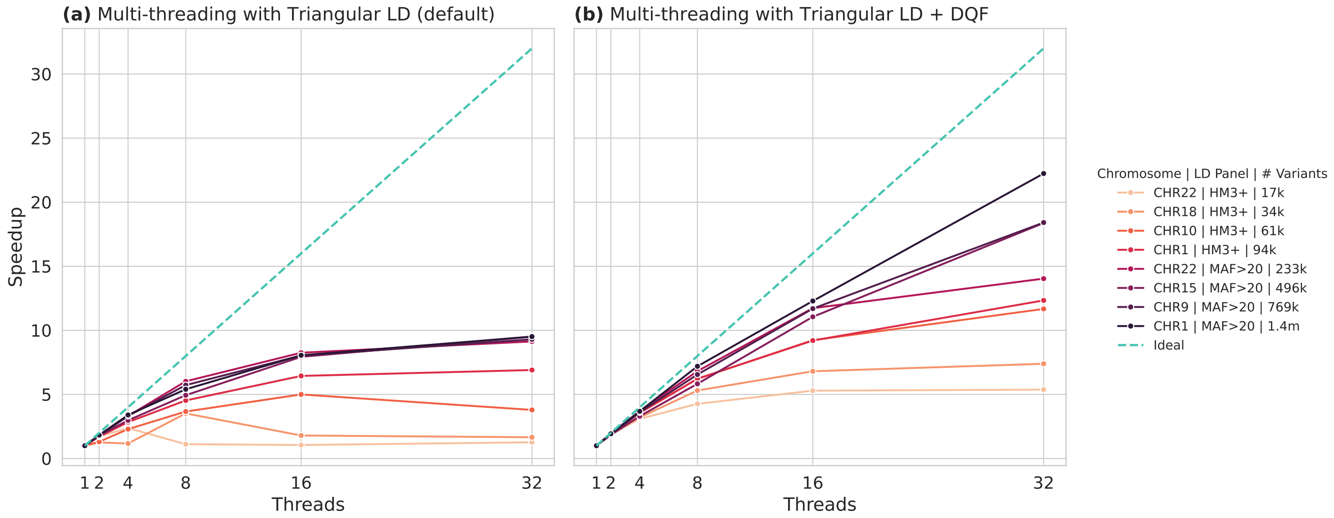


Figure S4: Speedup in runtime-per-iteration as a function of the number of threads across chromosomes of different sizes and two variants sets. Panel **(a)** shows speed up when using default LD mode (Triangular LD) and Panel **(b)** shows scaling when using Triangular LD + dequantize-on-the-fly (DQF) option. Each bold line corresponds to a combination of a chromosome (e.g. CHR1 is Chromosome 1) and variant set: HM3+ (HapMap3+) and MAC>20 (18m). Darker colors denote larger and denser chromosomes, where we expect to see more benefit from multi-threading. Dashed-line shows the ideal speedup expected if the runtime-per-iteration scales linearly with the number of threads.

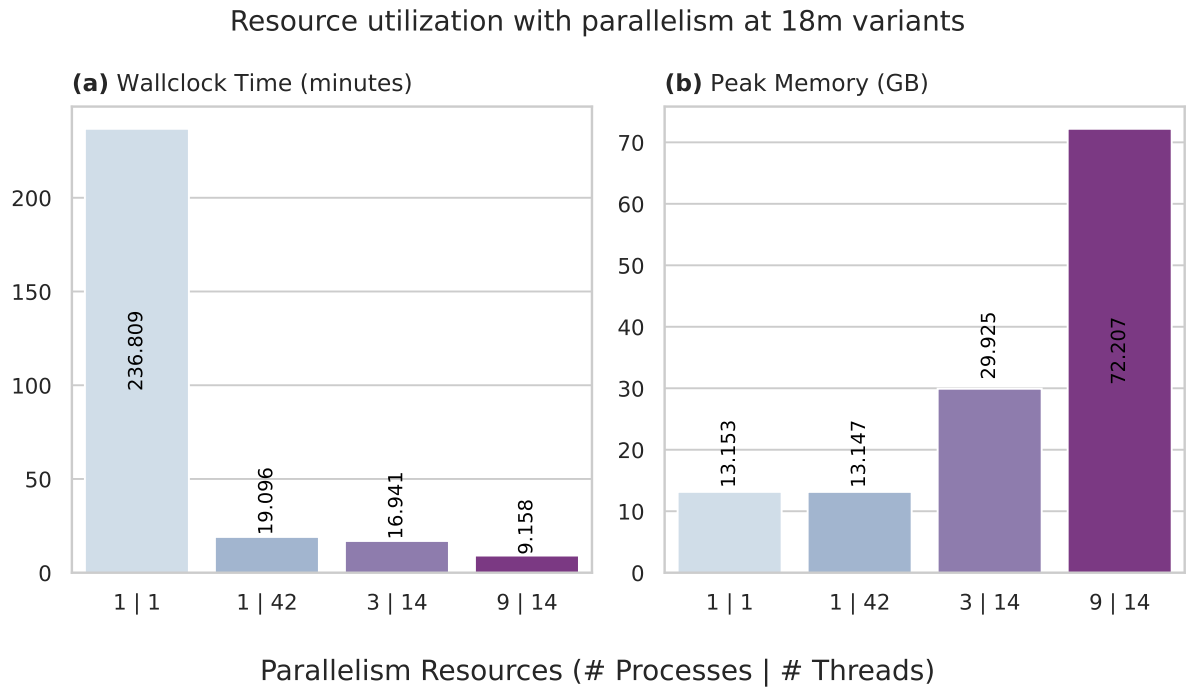


Figure S5: Resource utilization as a function of the number of cores and parallelism strategy (processes vs. threads) when performing inference using the MAC>20 (18 million) variant set. The experiments were conducted using GWAS summary statistics for Standing Height. Panel **(a)** shows total wallclock time as a function of resource configuration (number of processes vs. number of threads). Panel **(b)** shows peak memory (GB) as a function of the same configurations. Multi-threading improves total wallclock time without significantly affecting memory utilization.

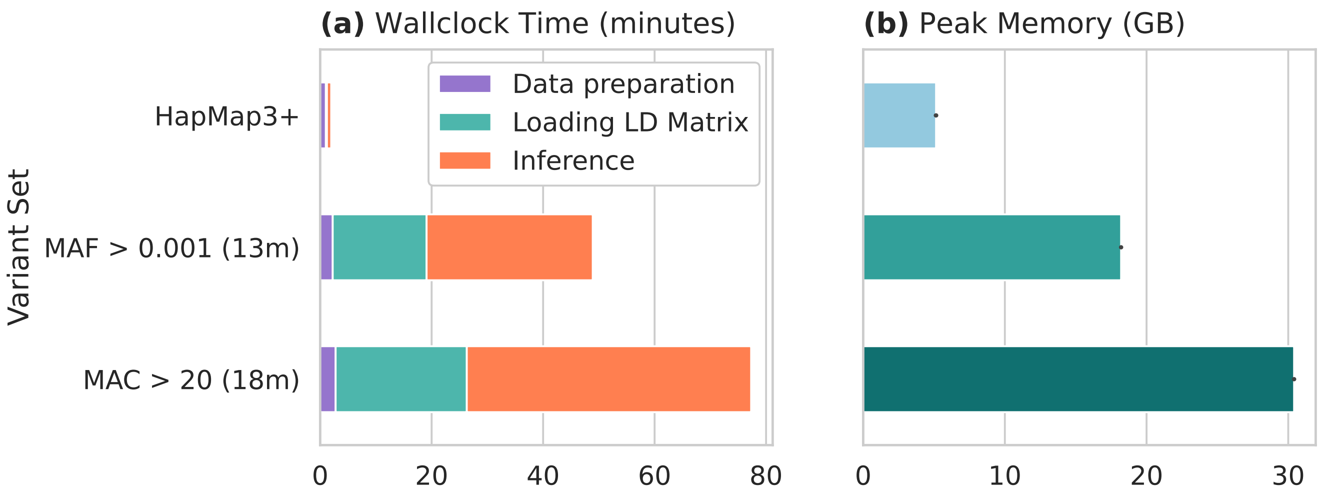


Figure S6: Computational characteristics of performing PRS inference using `VIPRS v0.1` on 75 continuous phenotypes in the Pan-UKB and across three variant sets. Unlike the main text, here we use LD matrices stored using `int16` quantization. Panels **(a-b)** show the average wallclock time (minutes) and peak memory usage (GB) across all the phenotypes. Colors in panel **(a)** denote average time required for each task during inference. Total wallclock time is mainly slowed due to increased time to fetch and filter the LD data. As expected, memory utilization is roughly twice what we obtained with `int8` quantization.

(a) Minimum eigenvalue of windowed LD matrices for European samples ($N = 362446$) in the UK Biobank

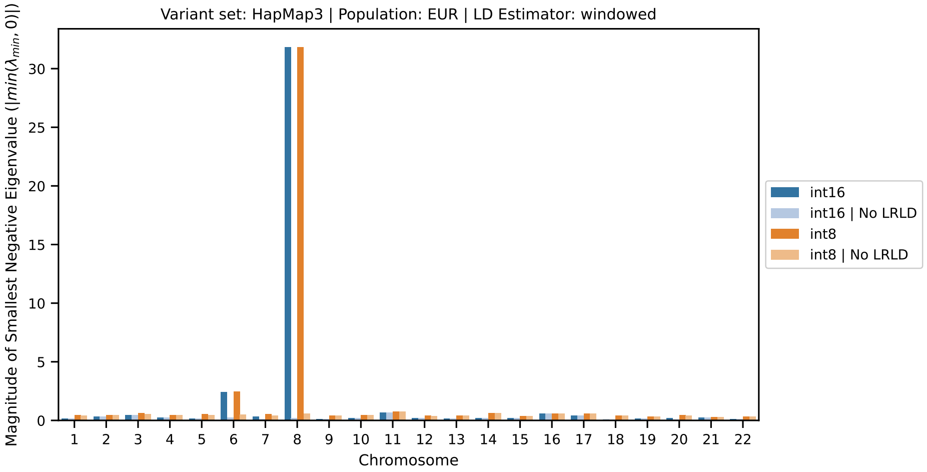


(b) Minimum eigenvalue of windowed LD matrices for East Asian samples ($N = 2700$) in the UK Biobank

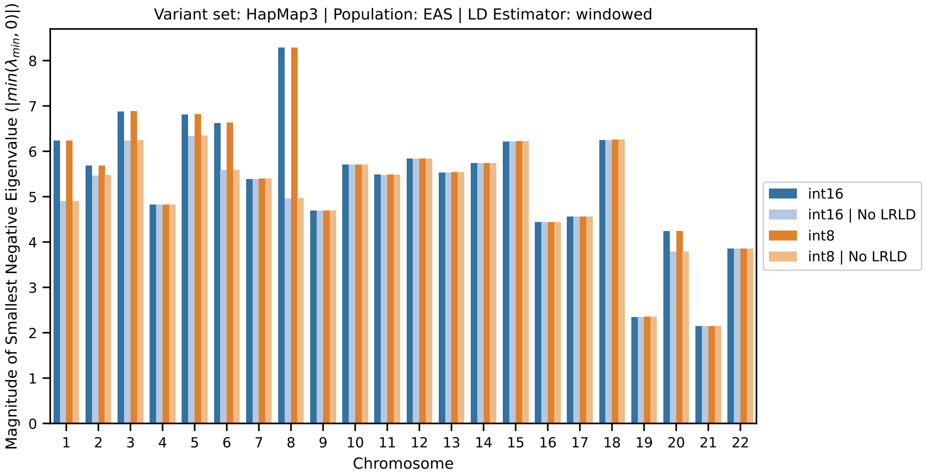


(c) Minimum eigenvalue of windowed LD matrices for African samples ($N = 6255$) in the UK Biobank

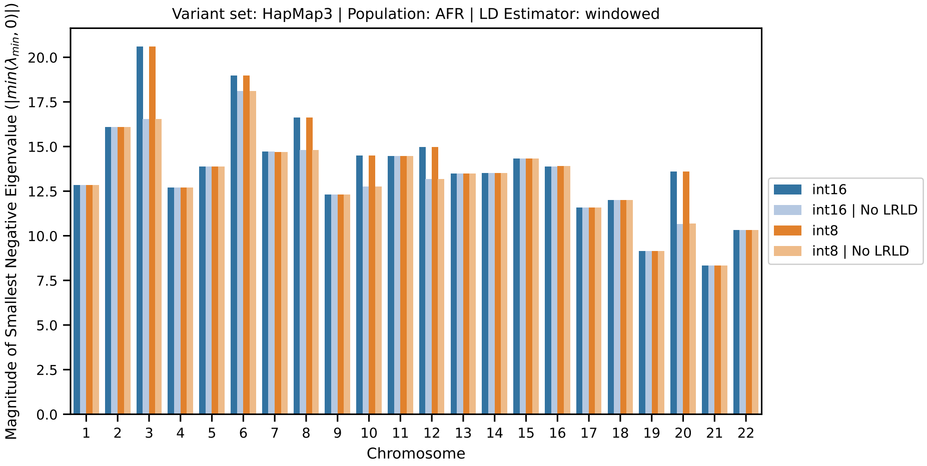


Figure S7: Absolute value of minimum (negative) eigenvalue for windowed (i.e. banded) LD matrices computed for three populations in the UK Biobank. The LD matrices were computed with window size of 3 centi Morgan. Colors denote data type used to store LD matrices: `int16` (blue) and `int8` (orange). Color shades denote LD matrices with (dark) and without (light) variants in long-range LD regions.

(a) Minimum eigenvalues for block-diagonal matrices using HapMap3+ variant set

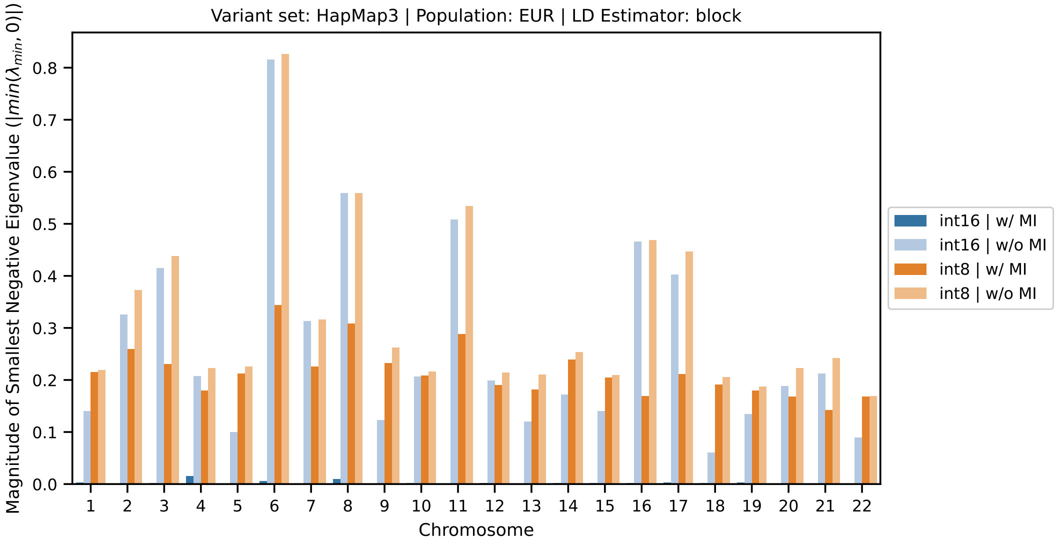


(b) Minimum eigenvalues for block-diagonal matrices using MAF > 0.001 (13m) variant set

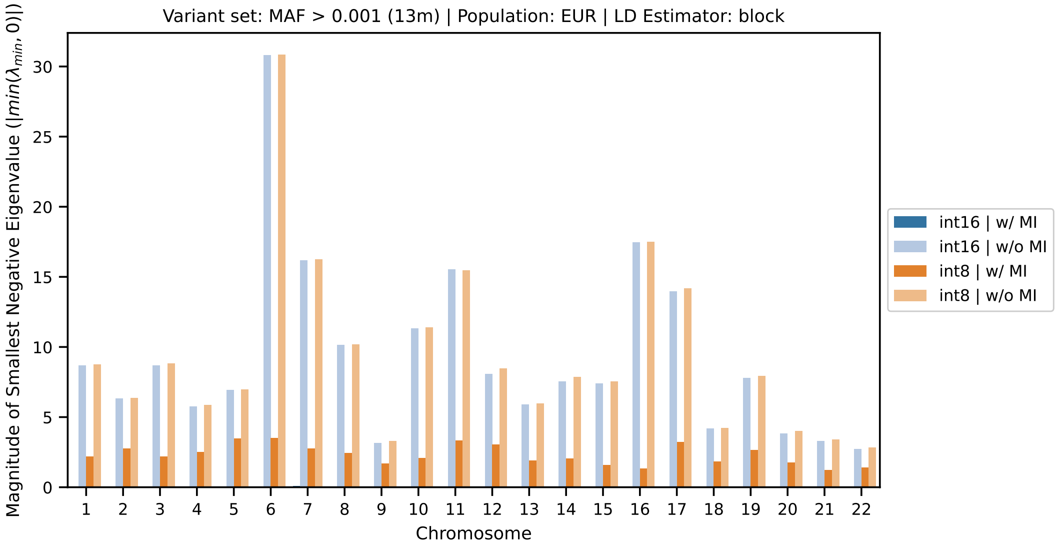


Figure S8: Absolute value of minimum (negative) eigenvalue for block-diagonal LD matrices computed for European samples in the UK Biobank across two variant sets: HapMap3+ (1.4m) and MAF>0.001 (13m). Colors denote data type used to store LD matrices: `int16` (blue) and `int8` (orange). Color shades denote whether Mean Imputation (MI; dark) was used to impute missing genotypes or missing observations were discarded (light).

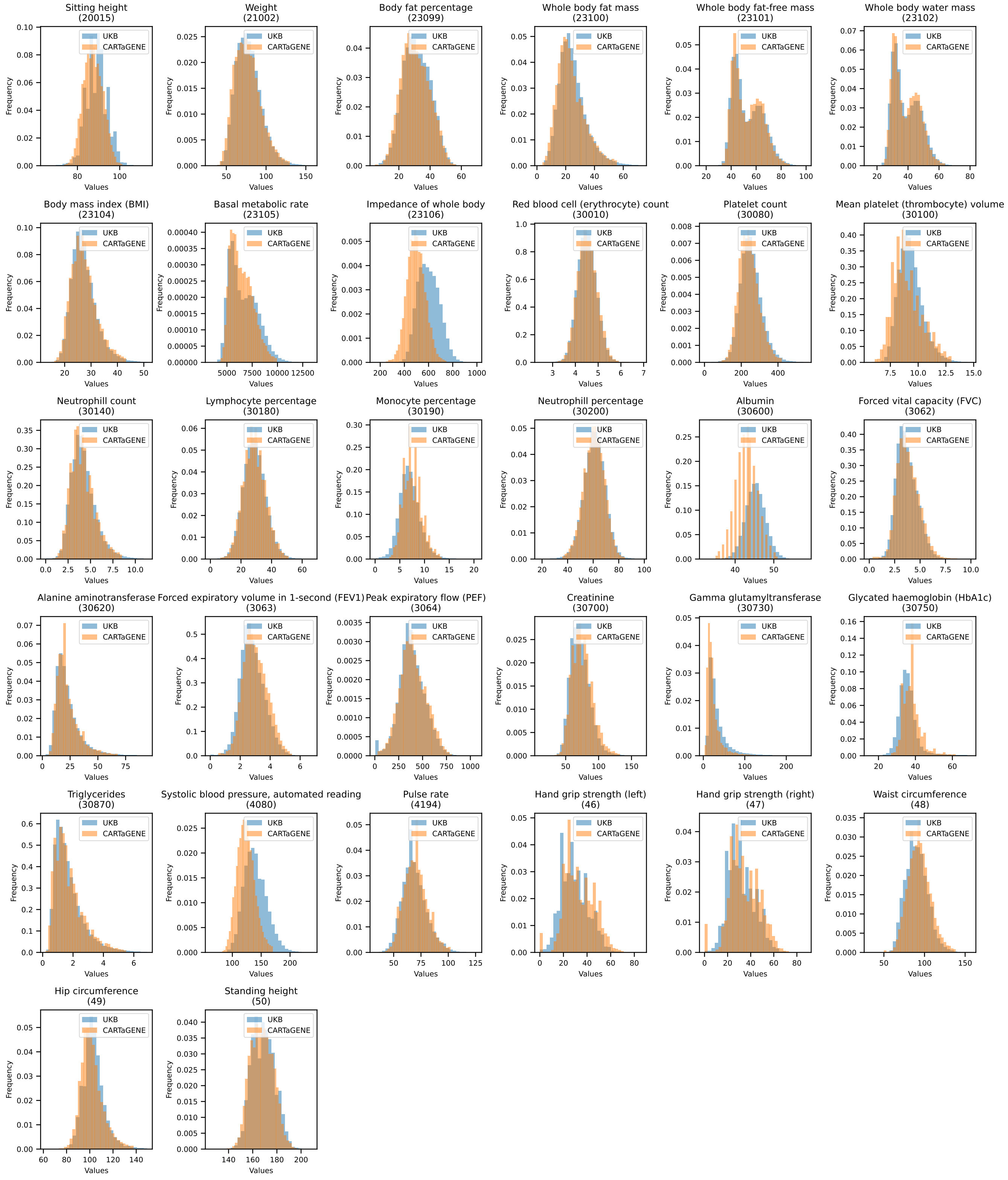


Figure S9: Cross-biobank phenotype distribution comparison between samples in the UK Biobank and the CARTaGENE cohort (Quebec, Canada) across the 32 of the 75 most heritable phenotypes in the Pan-UKB resource. Each panel shows the distribution for a particular phenotype and colored histograms show distribution in UKB (blue) and CARTaGENE (orange) samples.

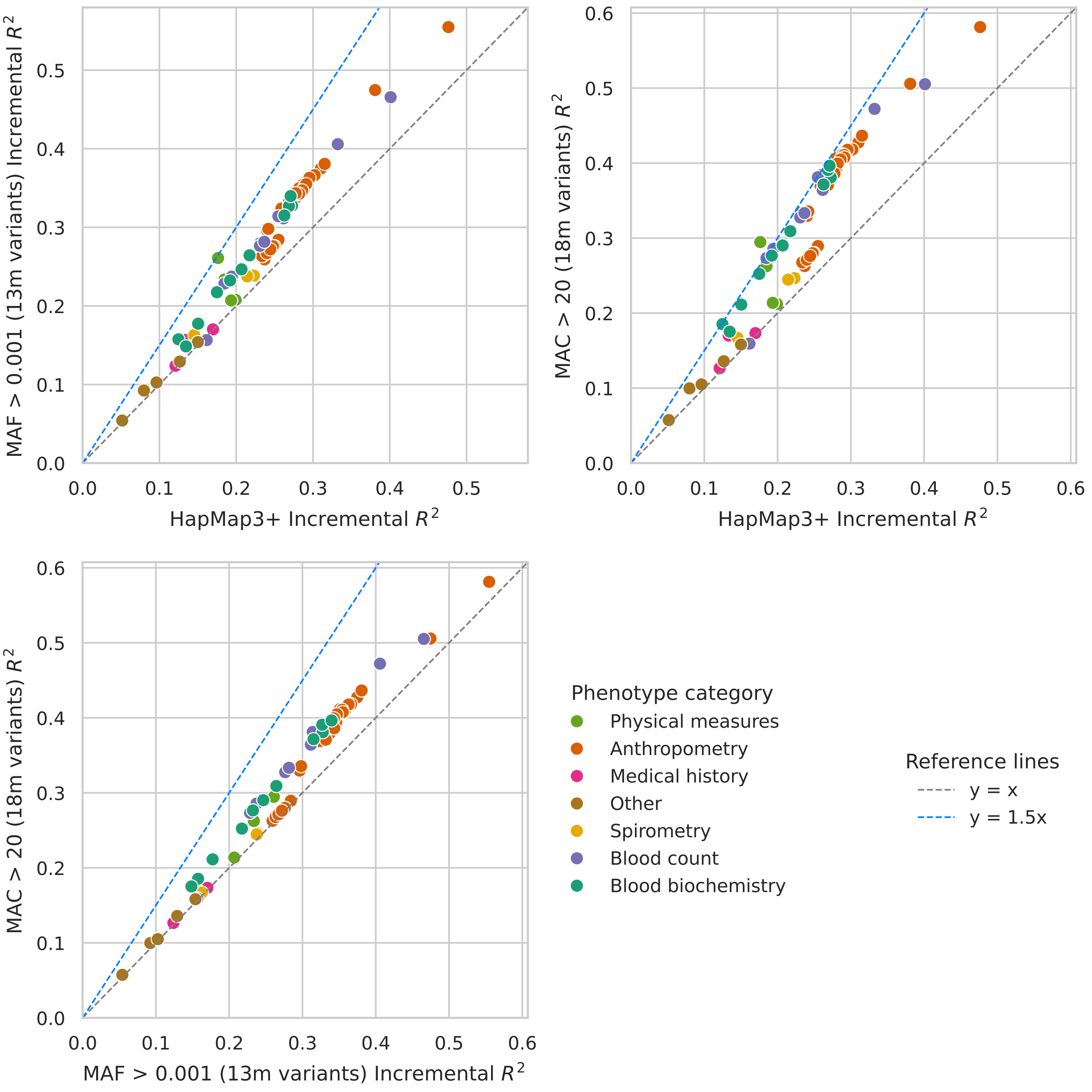


Figure S10: Comparison of prediction accuracy in the training cohort (EUR) in the Pan-UKB data resource using three variant sets: HapMap3+, MAF > 0.001 (13m), and MAC > 20 (18m). In each panel, we compare prediction accuracy (incremental $R^2$) on European samples in the UK Biobank when training PRS models using two of the three variant sets. Each dot shows prediction accuracy for one of the 75 phenotypes and colors denote the phenotype category.

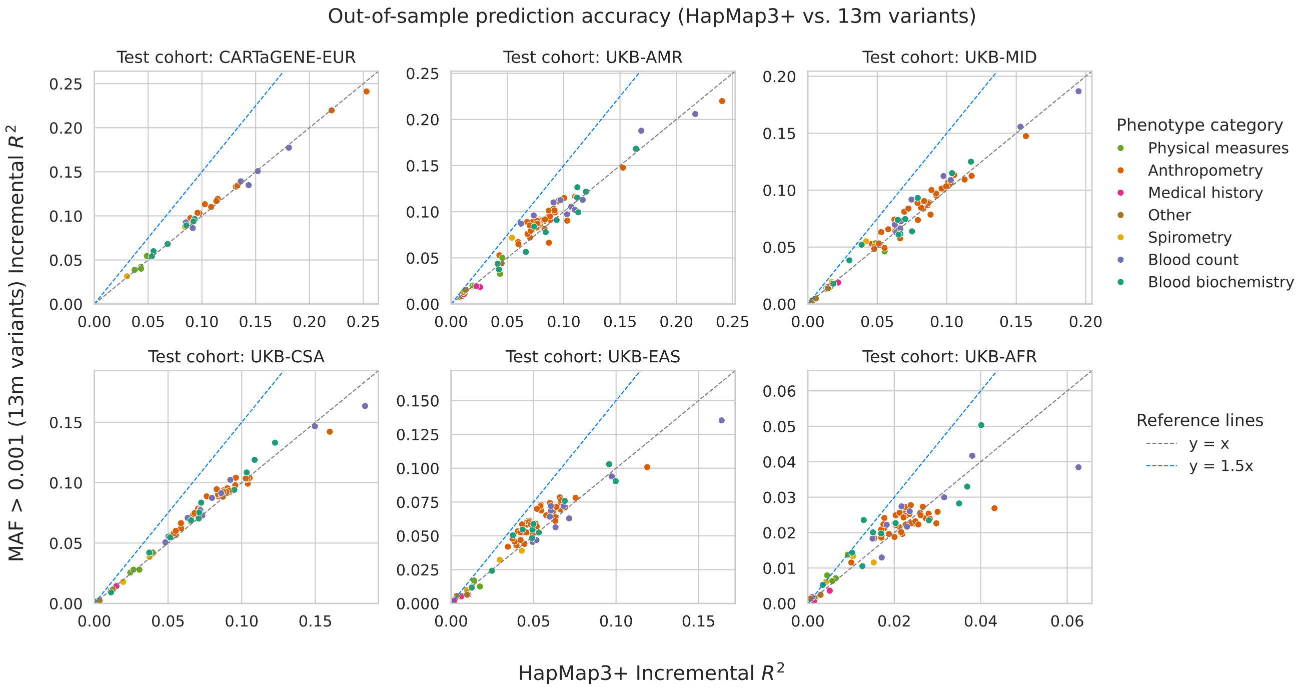


Figure S11: Systematic evaluation of PanUKB-derived PRS models across 75 continuous phenotypes and two variant sets. All polygenic scores were inferred from European GWAS summary statistics from the PanUKB initiative using `VIPRS v0.1`. The figure shows the comparative prediction accuracy between the HapMap3+ variant set on the x-axis (1.4 million variants) compared to the MAF>0.001 (13m) variant set on the y-axis. Each sub-panel compares the prediction accuracy for one of six held-out test cohorts across two biobanks: CARTaGENE and UK Biobank (UKB). The ancestry groups are EUR (European), AMR (Admixed American), MID (Middle Eastern), CSA (Central and South Asian), EAS (East Asian), and AFR (African). Colors denote different phenotype categories and dashed lines delineate the magnitude of the improvement in prediction accuracy.

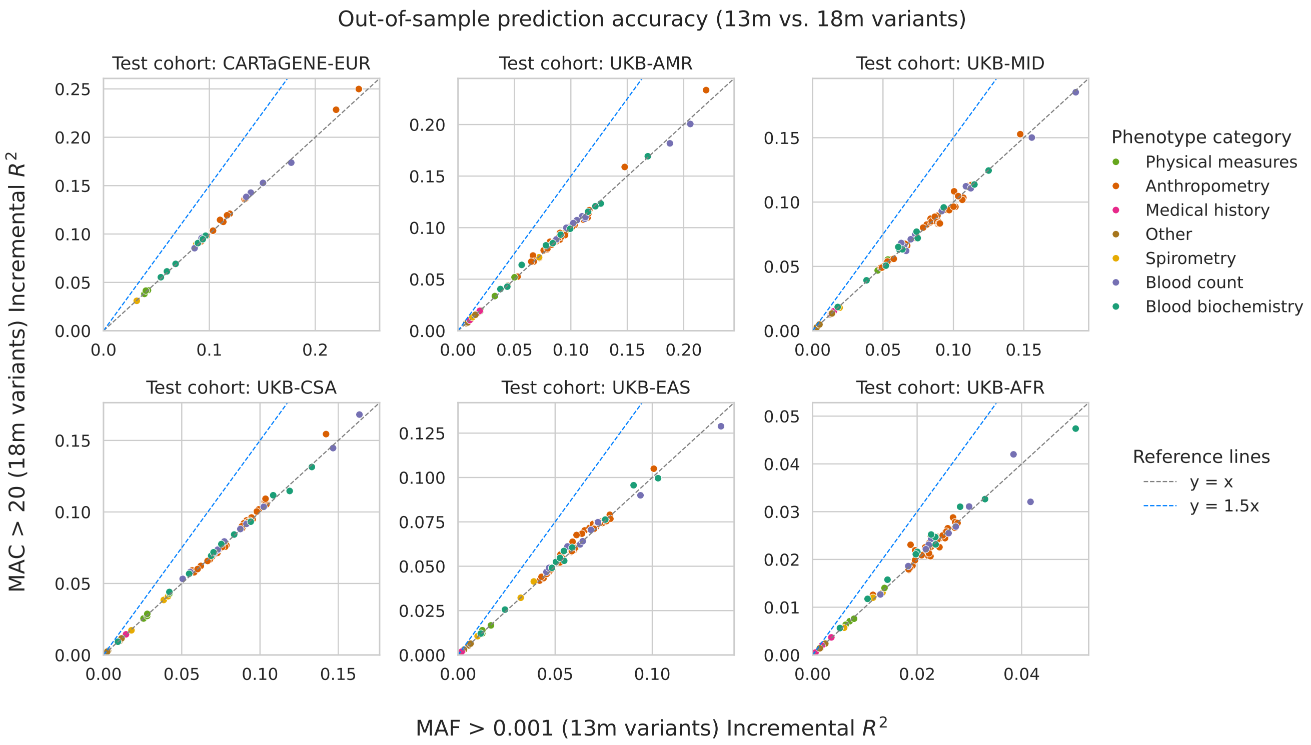


Figure S12: Systematic evaluation of PanUKB-derived PRS models across 75 continuous phenotypes and two variant sets. All polygenic scores were inferred from European GWAS summary statistics from the PanUKB initiative using `VIPRS v0.1`. The figure shows the comparative prediction accuracy between the MAF>0.001 (13m) variant set on the x-axis compared to the MAC>20 (18m) variant set on the y-axis. Each sub-panel compares the prediction accuracy for one of six held-out test cohorts across two biobanks: CARTaGENE and UK Biobank (UKB). The ancestry groups are EUR (European), AMR (Admixed American), MID (Middle Eastern), CSA (Central and South Asian), EAS (East Asian), and AFR (African). Colors denote different phenotype categories and dashed lines delineate the magnitude of the improvement in prediction accuracy.

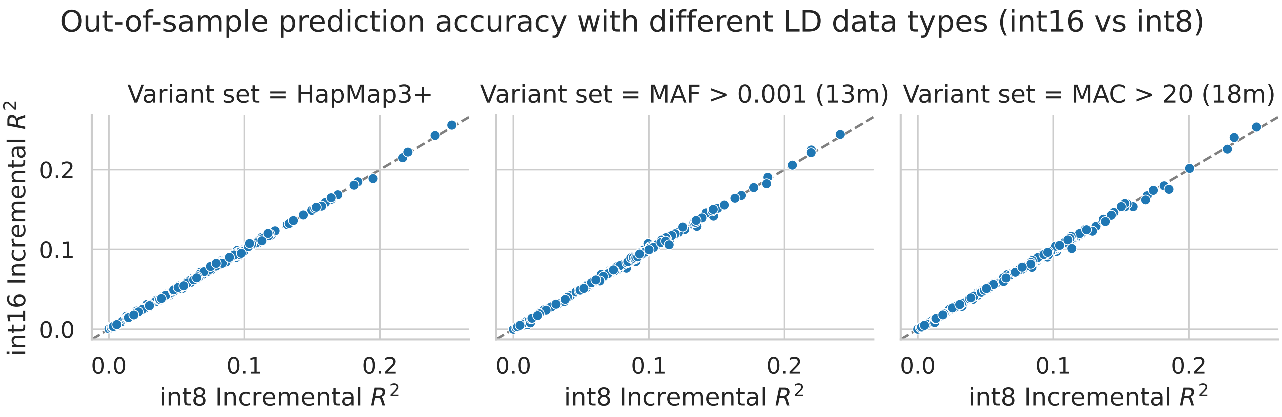


Figure S13: Comparison of held-out test prediction accuracy obtained when using LD matrices with `int8` (x-axis) vs. `int16` (y-axis) quantization across 75 phenotypes and six cohorts. All polygenic scores were inferred from European GWAS summary statistics from the PanUKB initiative using `VIPRS v0.1`. The held out cohorts include European samples from CARTaGENE as well as five non-European ancestry groups from the UK Biobank. This figure demonstrates that quantization to `int8` does not meaningfully affect prediction accuracy in most cases.

#### S2 Supplementary Tables

Table S1: Pan-UK Biobank phenotypes analyzed in this study. The table includes the UKB phenotype code, phenotype description, general category, as well as LDSC heritability estimates in European samples (provided by Pan-UKB manifest [1]).

| Phenocode | Description | Category | LDSC $h_2$ (EUR) |
|---|---|---|---|
| 48 | Waist circumference | Anthropometry | 0.223 |
| 49 | Hip circumference | Anthropometry | 0.250 |
| 50 | Standing height | Anthropometry | 0.588 |
| 51 | Seated height | Anthropometry | 0.291 |
| 20015 | Sitting height | Anthropometry | 0.449 |
| 21001 | Body mass index (BMI) | Anthropometry | 0.262 |
| 21002 | Weight | Anthropometry | 0.300 |
| 23098 | Weight | Anthropometry | 0.296 |
| 23099 | Body fat percentage | Anthropometry | 0.252 |
| 23100 | Whole body fat mass | Anthropometry | 0.275 |
| 23101 | Whole body fat-free mass | Anthropometry | 0.348 |
| 23102 | Whole body water mass | Anthropometry | 0.342 |
| 23104 | Body mass index (BMI) | Anthropometry | 0.268 |
| 23105 | Basal metabolic rate | Anthropometry | 0.336 |
| 23106 | Impedance of whole body | Anthropometry | 0.278 |
| 23107 | Impedance of leg (right) | Anthropometry | 0.253 |
| 23108 | Impedance of leg (left) | Anthropometry | 0.261 |
| 23109 | Impedance of arm (right) | Anthropometry | 0.268 |
| 23110 | Impedance of arm (left) | Anthropometry | 0.253 |
| 23111 | Leg fat percentage (right) | Anthropometry | 0.244 |
| 23112 | Leg fat mass (right) | Anthropometry | 0.255 |
| 23113 | Leg fat-free mass (right) | Anthropometry | 0.319 |
| 23114 | Leg predicted mass (right) | Anthropometry | 0.323 |
| 23115 | Leg fat percentage (left) | Anthropometry | 0.242 |
| 23116 | Leg fat mass (left) | Anthropometry | 0.256 |
| 23117 | Leg fat-free mass (left) | Anthropometry | 0.326 |
| 23118 | Leg predicted mass (left) | Anthropometry | 0.316 |
| 23119 | Arm fat percentage (right) | Anthropometry | 0.240 |
| 23120 | Arm fat mass (right) | Anthropometry | 0.247 |
| 23121 | Arm fat-free mass (right) | Anthropometry | 0.308 |
| 23122 | Arm predicted mass (right) | Anthropometry | 0.313 |
| 23123 | Arm fat percentage (left) | Anthropometry | 0.245 |
| 23124 | Arm fat mass (left) | Anthropometry | 0.255 |

| | | | |
|---|---|---|---|
| 23125 | Arm fat-free mass (left) | Anthropometry | 0.303 |
| 23126 | Arm predicted mass (left) | Anthropometry | 0.305 |
| 23127 | Trunk fat percentage | Anthropometry | 0.242 |
| 23128 | Trunk fat mass | Anthropometry | 0.265 |
| 23129 | Trunk fat-free mass | Anthropometry | 0.339 |
| 23130 | Trunk predicted mass | Anthropometry | 0.339 |
| 30600 | Albumin | Blood biochemistry | 0.145 |
| 30610 | Alkaline phosphatase | Blood biochemistry | 0.205 |
| 30620 | Alanine aminotransferase | Blood biochemistry | 0.124 |
| 30630 | Apolipoprotein A | Blood biochemistry | 0.182 |
| 30700 | Creatinine | Blood biochemistry | 0.213 |
| 30720 | Cystatin C | Blood biochemistry | 0.230 |
| 30730 | Gamma glutamyltransferase | Blood biochemistry | 0.180 |
| 30750 | Glycated haemoglobin (HbA1c) | Blood biochemistry | 0.210 |
| 30770 | IGF-1 | Blood biochemistry | 0.252 |
| 30870 | Triglycerides | Blood biochemistry | 0.177 |
| 30890 | Vitamin D | Blood biochemistry | 0.071 |
| 30010 | Red blood cell (erythrocyte) count | Blood count | 0.214 |
| 30080 | Platelet count | Blood count | 0.282 |
| 30100 | Mean platelet (thrombocyte) volume | Blood count | 0.281 |
| 30140 | Neutrophill count | Blood count | 0.174 |
| 30180 | Lymphocyte percentage | Blood count | 0.157 |
| 30190 | Monocyte percentage | Blood count | 0.160 |
| 30200 | Neutrophill percentage | Blood count | 0.149 |
| 30250 | Reticulocyte count | Blood count | 0.208 |
| 30270 | Mean sphered cell volume | Blood count | 0.206 |
| 30300 | High light scatter reticulocyte count | Blood count | 0.225 |
| 135 | Number of self-reported non-cancer illnesses | Medical history | 0.069 |
| 2178 | Overall health rating | Medical history | 0.109 |
| 2217 | Age started wearing glasses or contact lenses | Medical history | 0.081 |
| 400 | Time to complete round | Other | 0.096 |
| 1180 | Morning/evening person (chronotype) | Other | 0.130 |
| 1239 | Current tobacco smoking | Other | 0.062 |
| 1717 | Skin colour | Other | 0.081 |
| 30530 | Sodium in urine | Other | 0.077 |
| 46 | Hand grip strength (left) | Physical measures | 0.122 |
| 47 | Hand grip strength (right) | Physical measures | 0.119 |
| 4080 | Systolic blood pressure, automated reading | Physical measures | 0.160 |
| 4194 | Pulse rate | Physical measures | 0.126 |
| 3062 | Forced vital capacity (FVC) | Spirometry | 0.223 |
| 3063 | Forced expiratory volume in 1-second (FEV1) | Spirometry | 0.205 |

| | | | |
|---|---|---|---|
| 3064 | Peak expiratory flow (PEF) | Spirometry | 0.112 |

| Ancestry group | Variant set | Number of variants | LD matrix storage size |
|---|---|---|---|
| AFR | HapMap3+ | $1\,258\,340$ | 0.77 GB |
| AMR | HapMap3+ | $1\,284\,038$ | 0.91 GB |
| CSA | HapMap3+ | $1\,378\,881$ | 0.72 GB |
| EAS | HapMap3+ | $1\,144\,401$ | 0.57 GB |
| MID | HapMap3+ | $1\,305\,098$ | 0.88 GB |
| EUR | HapMap3+ | $1\,431\,634$ | 0.63 GB |
| EUR | MAF $> 0.1\%$ | $13\,482\,709$ | 37.68 GB |
| EUR | MAC $> 20$ | $17\,708\,098$ | 59.54 GB |

Table S2: LD matrix storage size for each ancestry group in the Pan-UKB. For European samples, we show LD matrix storage size across three different variant sets. LD matrices were estimated using banded masks with 3 centiMorgan window size. Entries of the matrix are stored using `int8` quantization.

### References


[1] Konrad J. Karczewski, Rahul Gupta, Masahiro Kanai, Wenhan Lu, Kristin Tsuo, Ying Wang, Raymond K. Walters, Patrick Turley, Shawneequa Callier, Nikolas Baya, et al. "Pan-UK Biobank GWAS improves discovery, analysis of genetic architecture, and resolution into ancestry-enriched effects". In: *medRxiv* (2024). DOI: 10.1101/2024.03.13.24303864. eprint: https://www.medrxiv.org/content/early/2024/03/15/2024.03.13.24303864.full.pdf. URL: https://www.medrxiv.org/content/early/2024/03/15/2024.03.13.24303864.
